# Timing of integration into the chromosome is critical for the fitness of an integrative and conjugative element and its bacterial host

**DOI:** 10.1101/2022.11.14.516373

**Authors:** Saria A. McKeithen-Mead, Alan D. Grossman

## Abstract

Integrative and conjugative elements (ICEs) are major contributors to genome plasticity in bacteria. ICEs reside integrated in the chromosome of a host bacterium and are passively propagated during chromosome replication and cell division. When activated, ICEs excise from the chromosome and may be transferred through the ICE-encoded conjugation machinery into a recipient cell. Integration into the chromosome of the new host generates a stable transconjugant. Although integration into the chromosome of a new host is critical for the stable acquisition of ICEs, few studies have directly investigated the molecular events that occur in recipient cells during generation of a stable transconjugant. We found that integration of ICE*Bs1*, an ICE of *Bacillus subtilis*, occurred several generations after initial transfer to a new host. Premature integration in new hosts led to cell death and hence decreased fitness of the ICE and transconjugants. Host lethality due to premature integration was caused by rolling circle replication that initiated in the integrated ICE*Bs1* and extended into the host chromosome, resulting in catastrophic genome instability. Our results demonstrate that the timing of integration of an ICE is linked to cessation of autonomous replication of the ICE, and that perturbing this linkage leads to a decrease in ICE and host fitness due to a loss of viability of transconjugants. Linking integration to cessation of autonomous replication appears to be a conserved regulatory scheme for mobile genetic elements that both replicate and integrate into the chromosome of their host.

**Author Summary:** Horizontal gene transfer helps drive microbial evolution, enabling bacteria to rapidly acquire new genes and traits. Integrative and conjugative elements (ICEs) are mobile genetic elements that reside integrated in the chromosome of a host bacterium and can transfer to other cells via a contact-dependent mechanism called conjugation. Some ICEs contain genes that confer important traits to their bacterial hosts, including antibiotic resistances, metabolic capabilities, and pathogenicity. Central to the propagation of ICEs and other mobile genetic elements is the balance between dissemination to new hosts and maintenance within a host, while minimizing the fitness burden imposed on their hosts. We describe an underlying regulatory mechanism that allows for an ICE that has entered into a nascent host to replicate and potentially spread to other hosts before integration into the chromosome. Integration occurs shortly after or concomitant with cessation of ICE replication. Disruption of this regulatory link results in premature integration and fitness defects for both the host and the ICE; lethality for the host and reduced spread of the ICE.

## Introduction

Horizontal gene transfer (HGT) is a major driver of prokaryotic evolution, enabling the movement of genes from a donor organism to a recipient and subsequent maintenance of horizontally acquired genes within a population. Genes transferred horizontally can confer a wide range of traits, including antibiotic resistances, metabolic capacities, pathogenicity, and symbiosis (reviewed in [1]).Conjugation, a form of HGT, is the contact-dependent transfer of DNA from a donor to a recipient cell. Transfer is through a conjugation machine (a type 4 secretion system, T4SS) that is typically encoded by a conjugative element in a donor cell [2–4].

There are two broad classes of conjugative elements. Integrative and conjugative elements (ICEs, also known as conjugative transposons) normally reside integrated in a host genome and can become activated to excise and express the conjugation machinery [1,5]. Conjugative plasmids are extra-chromosomal and also contain genes needed for their transfer to recipient cells (1–3,6). Conjugative transfer of ICEs and plasmids originates at the origin of transfer (*oriT*) in the element and is initiated by the element-encoded relaxase protein that nicks and becomes covalently attached to the 5’-end of the nicked strand. The nicked DNA is unwound and linear single-stranded DNA (ssDNA) with the attached relaxase can be transferred from the donor to a recipient through the conjugation machinery [6]. Once inside the recipient, the ssDNA circularizes, and undergoes second-strand synthesis to generate circular dsDNA. ICE genes can now be expressed and the double-stranded circular form of the ICE is a substrate for integration into the host chromosome [1,5].

Unlike that for conjugative plasmids, stable acquisition of an ICE requires integration into the host chromosome where the ICE is then passively propagated as the host replicates and segregates its DNA to daughter cells. Many ICEs are known to undergo limited rounds of autonomous replication and both replication and integration are often essential for stable acquisition of the element [1,7–16]. The requirement for integration for stable acquisition of an ICE is a clear delineation between conjugative plasmids and ICEs and introduces an area for differential selection and evolution of regulatory schemes.

Little is known about the regulatory factors that drive integration and stable acquisition of an ICE after transfer to a new host. Analysis of events in transconjugants has typically relied on end-point readouts of overall mating efficiency to discern the importance of ICE functions, thus obscuring the context in which a particular function is important. A primary challenge for these types of studies is that transfer events are relatively infrequent and, for most ICEs, only a subpopulation of cells participate.

We used ICE*Bs1* (**Fig 1**) from *Bacillus subtilis* to investigate the timing of successful integration and acquisition in transconjugants. ICE*Bs1* is well-suited for investigating conserved ICE dynamics as it has been well-studied, is easily manipulated, and can be activated in a relatively large fraction (~20% - 90%) of cells in the population [17–19] ICE*Bs1* integrates via site-specific recombination into one site in the *B. subtilis* chromosome: in *trnS-leu2* which encodes a leucyl-tRNA [17,19]. The specific integration site (*attB*; <u>att</u>achment site in the <u>b</u>acterium) allows for precise measurements of integration (stable acquisition).

**Fig 1.**
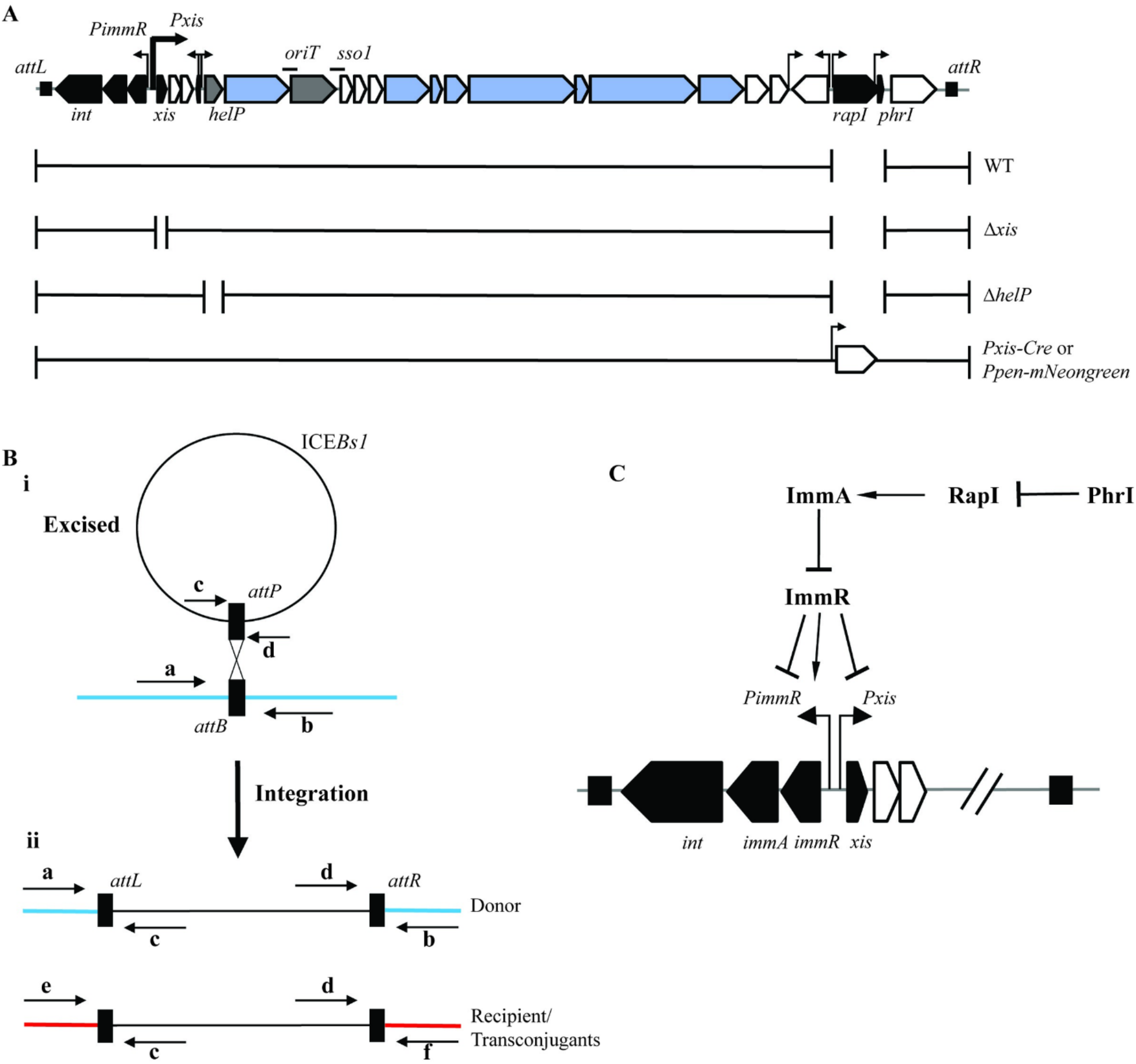
ICEBs1 integrated versus excised state per regulation with qPCR-assay schematic. **A)** Map of genes and some of the sites in ICE*Bs1*. Black rectangles at the end of ICE*Bs1* represent the flanking 60 bp repeats *attL* and *attR*, the junctions between the chromosome and ICE*Bs1* on the left and right, respectively. Genes are represented by horizontal block arrows, not precisely to scale, indicating the direction of transcription. Genes are colored according to function (black = regulation, white = cargo/unknown function, blue = conjugation, grey = replication). Black lines above ICE*Bs1* indicate *oriT*, the origin of transfer, and *sso1*, one of the single stranded origins of replication. Vertical right-angle arrows indicate promoters. Below the ICE*Bs1* gene map is a schematic of mutations. Gaps represent deletions (all strains have deletions of *rapI-phrI*). Insertions (P*xis*-*cre* and P*pen-mNeongreen*) are indicated by promoter arrows and horizontal block arrows. **B)** General schematic for detection of products of ICE*Bs1* excision and integration. PCR primers are indicated by arrows with letter a, b, c, d, e, or f. Different combinations of primers can detect different forms of ICE*Bs1*, as indicated. Black rectangles indicate ICE*Bs1* attachment sites: *attP* on ICE*Bs1*; *attB* on *B. subtilis* chromosome; *attL* and *attR* for the integrated element and the junctions with the chromosome.

i. Excised ICE in circular form with empty *attB* in the chromosome. ICE*Bs1* is represented by a black circle. The *B. subtilis* chromosome is represented by a blue line.
ii. Integrated ICE*Bs1* in original donors (top) and recipient/transconjugants (bottom). The different flanking sequences in the experimental set-up are shown (blue for donors; red for recipients/transconjugants). PCR primers a and b are specific for the ectopic locus (*amyE*::*attB*) in donors, and primers e and f are specific for the native locus in recipients and transconjugants. For example, the combination of primers d and f would detect integration of ICE*Bs1* (*attR*) at its native site *attB*. **C)ICE*Bs1* regulation.** The ICE*Bs1* gene products RapI (activator), ImmA (anti-repressor and protease), ImmR (repressor) all regulate ICE*Bs1* gene expression from the major promoter P*xis*. P*xis* is repressed by ImmR. When RapI is present and active, it causes the protease ImmA to cleave ImmR, thereby causing de-repression of transcription from P*xis* [21,22]. ImmR both activates and represses transcription from its own promoter (P*immR*) [18].

Using a combination of population-based and single-cell assays, we found that integration in transconjugants occurs several generations after transfer of the element. Deletion of *xis* (encoding the recombination directionality factor that is needed for element excision from the chromosome) caused premature integration in transconjugants and was lethal for most transconjugants. Lethality was caused by integration of an active element that was undergoing autonomous rolling circle replication. Our results highlight the importance of ICE regulation in transconjugants and reveal the important underlying linkage between the cessation of autonomous replication (a plasmid-like function) and integration.

## Results

### Timing of ICE*Bs1* transfer and integration in transconjugants

Integration of an ICE into the genome of a new host is essential for the stable acquisition of the element. Following the transfer of ICE*Bs1* from a donor to recipient, the element is capable of rapidly transferring through a chain of cells, indicating that the element is active for a period of time before it integrates into the chromosome [20].We sought to determine when ICE*Bs1* integrates into the chromosome of transconjugants.

#### Strains for distinguishing integration in donors and transconjugants

ICE*Bs1* has a unique integration site (*attB*) in the bacterial chromosome that is normally identical in both donor and recipient cells, making it virtually impossible to physically distinguish, at the molecular level, integration in transconjugants from that in the initial donor. In order to detect integration in transconjugants separately from the initial donors, we constructed a donor strain (SAM078) in which the site of ICE*Bs1* integration was in *amyE* (a nonessential gene often used for inserting cloned genes) rather than its normal position in *trnS-leu2*. We could then distinguish integrants in the transconjugant population (integration in the normal chromosomal site) from those in the initial donor (at *amyE*, the ectopic location) using PCR primer pairs that were specific for transconjugants (**Fig 1B**). We normalized the number of integrants in the transconjugant population relative to a genomic marker (*mls*, that confers resistance to erythromycin and lincomycin) that was not present in donors.

#### Timing of acquisition of ICE*Bs1*

We measured the conjugation efficiency at various times after the start of mating (Methods). Transconjugants were determined by the number of colony forming units (CFUs) with indicative antibiotic resistances (Methods). Recipients (SAM103) lacked ICE*Bs1* (ICE*Bs1*^0^). ICE*Bs1* in donors (SAM078) was activated by overexpression of *rapI* from an inducible promoter at an ectopic locus (Methods). RapI causes the ICE*Bs1*-encoded protease ImmA to cleave the element repressor ImmR, thereby causing de-repression of ICE*Bs1* (**Fig 1C**) [17,18,21,22]. After 2 hours, there were typically 5-10% transconjugants per initial donor (**Table 1**). After 3 hours, the frequency increased to approximately 30% transconjugants per donor (**Table 1**). The detection of transconjugants as CFUs at any time post-mating indicated that ICE*Bs1* had transferred from a donor to a recipient, but did not indicate when ICE*Bs1* had integrated into the chromosome of the transconjugant.

**Table 1.** Effect of *xis* on ICE*Bs1* mating^1^.

| Time | Donors | Mating efficiency (%)<sup>2</sup> |  |
| --- | --- | --- | --- |
|  |  | Recipients |  |
|  |  | WT (SAM103) | P<sub>xis</sub>-*xis* (SAM379) |
| 2 hrs | WT (SAM078) | 9.5 ± 1.6 | 7.4 ± 0.2 |
| | Δ*xis* P<sub>spank</sub>-*xis* (SAM207) | 0.58 ± 0.3 | 1.3 ± 0.5 |
| 3 hrs | WT (SAM078) | 30.7 ± 2.9 | 25.8 ± 0.9 |
| | Δ*xis* P<sub>spank</sub>-*xis* (SAM207) | 16.7 ± 1.9 | 32.7 ± 6.5 |

<sup>1</sup> Mating experiments were performed on cellulose nitrate filters on agar plates as described in Methods. Expression of *rapI* was induced by addition of 1% xylose and 1% arabinose to donor cultures 40 mins before mixing donor and recipients cells. All recipients are cured of ICEBsI (ICEBsI<sup>0</sup>) and contain *mls* (erythromycin resistant). Strain numbers are in parentheses. All donor strains contain ICEBsI $\Delta$ (*rapI*-*phrI342*):*kan* (kanamycin resistant) and P<sub>xyl</sub>-*rapI*.
<sup>2</sup> Mating efficiency is indicated by the percentage of CFUs of transconjugants per initial donor. Efficiencies are the average of at least three independent experiments ± standard error of the mean.

#### Timing of ICE*Bs1* integration in transconjugants

We measured the fraction of transconjugants in which ICE*Bs1* had integrated at two and three hours after the start of mating. Briefly, genomic DNA was isolated from the cells that were recovered from mating filters and we used qPCR to measure integration of ICE*Bs1*. Because the location of the integration site was different in donors (SAM078) and recipients (SAM103), we could measure integration of ICE*Bs1* in the transconjugants (recipients) using a primer pair that would detect the junction between the right end of ICE*Bs1* and the host chromosome (*attR*) that is created only if ICE*Bs1* had integrated into the chromosome of a recipient. We also determined the total number of recipients in the population by qPCR of a sequence (*mls*) present in recipients (and transconjugants), but not in donors.

We found that ICE*Bs1* had integrated in approximately 5% of the transconjugants after two hours and approximately 70% after three hours of mating (**Fig 2A**). That is, of the ~10% transconjugants per donor after two hours of mating (Table 1), only 5% of that ~10% contained an integrated ICE*Bs1*. Of the ~30% transconjugants per donor after three hours of mating (Table 1), ~70% contained an integrated ICE*Bs1*. Based on these findings, we conclude that there was a delay between the time of ICE*Bs1* transfer and integration and that by 3 hours post-mating, ICE*Bs1* had integrated into the chromosome in the majority of transconjugants.

**Fig 2.**
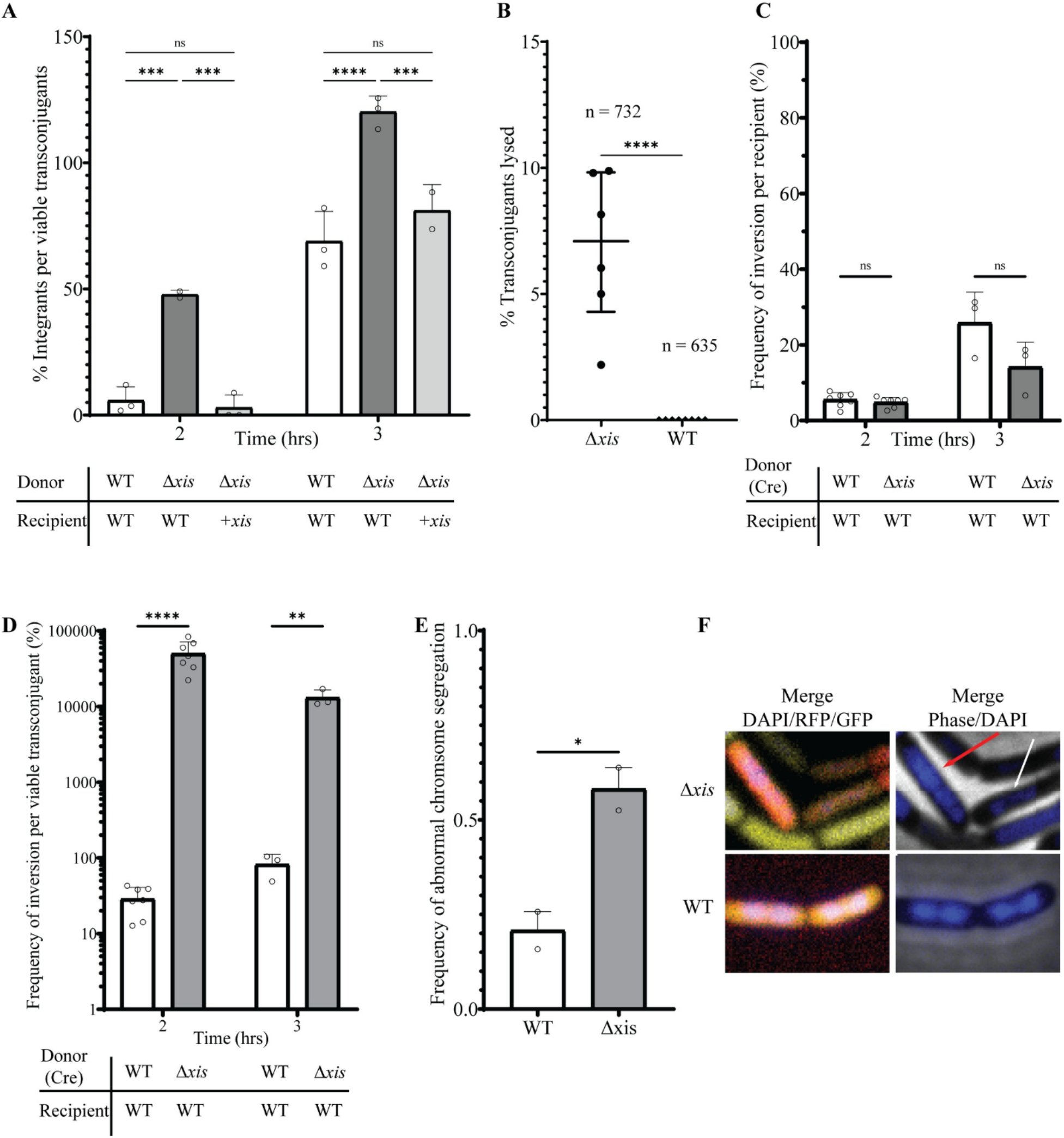
Early integration of ICE*Bs1* causes cell death. **A)** Integration of ICE*Bs1* in transconjugants was measured by qPCR and normalized to the number of transconjugant CFUs. The identity of the donor and recipient strains is indicated below the x-axis. Donors: WT (SAM078); Δ*xis* (SAM207). Recipients: WT (SAM103); +*xis* (*xis* expressed in recipients; SAM379). Mating pairs are shown below the graph with relevant genotypes indicated. Circles indicate the result from independent experiments. Error bars indicate standard deviation. Statistical comparison: two-way Anova, Tukey’s multiple comparison test: *** P value < 0.005; **** P value < 0.0001. **B)** Lysis of ICE*Bs1* Δ*xis* transconjugants. Donors (ICE*Bs1* Δ*xis;* SAM472; or ICE*Bs1 xis+*; SAM288) harboring ICE*Bs1* with constitutively expressed *mNeongreen* were mated with recipients (SAM271) constitutively expressing *mApple* (and MLS resistant) for 4 hr. Samples were then spotted onto nutrient-rich agarose pads containing erythromycin and lincomycin (selecting for recipients and transconjugants, preventing growth of donors). Transconjugants were identified by co-expression of *mNeongreen* and *mApple*. Cell fate was tracked over the course of 3 hr. Each point represents the frequency of lysed transconjugant cells in an individual field of cells from one representative experiment. Top and bottom lines indicate the interquartile range and the middle line indicates the mean. We tracked 732 and 635 transconjugants from Δ*xis* and wild type (*xis*+), respectively. Of these 51 (~7%) of the Δ*xis* and 1 (≤0.2%) of the *xis*+ transconjugants lysed during the 4 hrs of observation. Statistical comparison: two-tailed t-test, **** P value <0.0001. **C)** Acquisition of ICE*Bs1* as measured by Cre-mediated recombination per recipient. Acquisition efficiency was measured by qPCR by amplifying across the inversion sequence (which would indicate Cre-mediated recombination) and normalizing to a unique sequence (*mls*) in recipients. Donors: WT (SAM830); Δ*xis* donor (SAM892). Recipients: WT (SAM599). Error bars indicate standard deviation. Statistical comparison: t-test, Holm-Šídák method, ns indicates not significant. **D)** Same experiment as in Fig 2C. qPCR from amplification across inversion sequence normalized to transconjugant CFUs. These are the data from Fig 2C divided by data from corresponding strains and conditions from S2 Fig. The frequency of inversions per viable transconjugant for Δ*xis* decreased at 3 hr because the number of viable transconjugants increased. Statistical comparison: t-test, Holm-Šídák method, ** P value < 0.005; **** P value < 0.00001. **E)** Experimental set-up as in **B** with the addition of DAPI to agarose pads. Chromosomal abnormalities were tracked over the course of 3 h. Results are from randomly sampling between 100 – 116 cells for each condition. Data were analyzed by two individuals who did not know the identity of the samples. Error bars indicate standard error of the mean. Statistical comparison: two-tailed t test, * P value < 0.05. **F)** Representative micrograph images showing abnormal (Δ*xis*) and normal (WT) chromosome segregation in transconjugants from the experiment in Fig 2E. Red arrow indicates cell with an elongated nucleoid. White arrow indicates a cell with asymmetrical chromosome segregation.

### Loss of *xis* caused premature integration in transconjugants

In many integrative elements, including temperate phages and ICEs, recombination into and out of the host chromosome is catalyzed by a recombinase, often named Int (integrase). For many elements, a recombination directionality factor, often called Xis or Rdf (excisionase or recombination directionality factor) is needed, along with Int, for recombination (excision) out of the genome. ICE*Bs1* encodes both Int and Xis; Int is needed for both integration and excision, and Xis is needed just for excision [19]. Regulation of ICE*Bs1* is such that *int* is expressed at a low level constitutively, and *xis* is expressed at a high level immediately upon de-repression of ICE*Bs1* gene expression (**Fig 1A and 1C**) [19].

We found that loss of *xis* caused early integration in transconjugants. We used donor cells that contained ICE*Bs1* without *xis* (ICE*Bs1* Δ*xis*; SAM207). To enable excision, we provided *xis in trans* from an ectopic locus, essentially as described previously [19]. Thus, ICE*Bs1* Δ*xis* could excise in this donor but there would be no *xis* in transconjugants. At two hours post-mating, ICE*Bs1* Δ*xis* had integrated in ~50% of transconjugants, an increase of ~10-fold relative to that of wild-type ICE*Bs1* (*xis+*) (**Fig 2A**). Loss of *xis* also caused a decrease in the number of viable transconjugants per donor. At two hours post-mating, there were ~0.6% transconjugants per initial donor compared to ~10% for wild type (*xis+*) (**Table 1**).

Early integration of ICE*Bs1* Δ*xis* was due to the absence of *xis* in transconjugants and not loss of a site or an effect on a downstream gene. We expressed *xis* from its own promoter (P*xis*-*xis*, integrated as a single copy in the chromosome) in recipients. When these recipients (SAM379) were used in a mating with ICE*Bs1* Δ*xis* donors (SAM207), the timing of integration was similar to that of wild-type ICE*Bs1* (*xis*+) (**Fig 2A**). Additionally, conjugation efficiency in the recipient expressing *xis* increased relative to that of ICE*Bs1* Δ*xis* (**Table 1**). These results indicate that early integration of the *Δxis* mutant in transconjugants was due to loss of *xis* and not an unexpected secondary effect on downstream genes or loss of a site. We conclude that *xis* is needed in transconjugants for proper timing of integration.

### Early integration caused cell death

Our data indicated that many of the initial transconjugants that acquired ICE*Bs1* Δ*xis* were likely dying. For example, three hours post-mating, the integration efficiency per viable transconjugant of ICE*Bs1* Δ*xis* was greater than 100% (**Fig 2A**). This indicated that ICE*Bs1* Δ*xis* was integrating in the transconjugants, but that many of these transconjugants were unable to form colonies. The resulting increase in the ratio of integrants per transconjugant CFU was likely due to the decrease in transconjugant CFUs. To further investigate this, we monitored the fate of individual transconjugants using fluorescence microscopy.

To distinguish donors, recipients, and transconjugants, the three cell types present in a mating mix, we constructed recipients that constitutively produced red fluorescent protein from *mApple* and donors that constitutively produced yellow-green fluorescent protein from *mNeongreen* that had been introduced into ICE*Bs1* (ICE*Bs1 mNeongreen*). Transconjugants produce both mApple and mNeongreen and were easily distinguished from both donors and recipients that expressed only one fluorescent protein.

We found that a higher proportion of transconjugants that had acquired ICE*Bs1* Δ*xis* lysed compared to those that had acquired ICE*Bs1 xis+*. In these experiments, we mated donors (*xis*+, SAM288; Δ*xis*, SAM472) with recipients (SAM271) for four hours and spotted the mating mixture onto agarose pads to observe individual cells by time-lapse microscopy for three hours (Methods). Transconjugants were identified as cells that had both red (mApple) and yellow (mNeongreen) fluorescence. During the three hours of observation, ~7% (51 of 732 cells observed) of Δ*xis* transconjugants lysed, compared to less than 0.2% (1 of 635 cells observed) of wild-type (*xis*^+^) transconjugants (**Fig 2B, S1 Movie**). We suspect that Δ*xis* transconjugants continued to lose viability after the three hours of observation.

### Using the recombinase Cre to detect transfer of ICE*Bs1*, irrespective of viability of transconjugants

To better measure integration-mediated death of transconjugants, separate from the ability of transconjugants to form colonies, we developed a reporter system that uses Cre-mediated recombination to demonstrate transfer of ICE*Bs1*. We reasoned that if ICE*Bs1* Δ*xis* was integrating prematurely, then ICE*Bs1* genes must be expressed. If the element encoded a heterologous recombinase (e.g., Cre), we could detect transconjugants by virtue of a Cre-catalyzed recombination event, even if the transconjugants lost viability. Briefly, we inserted *cre* driven by P*xis* in place of *rapI-phrI* in ICE*Bs1* (*xis*+, SAM830; Δ*xis*, SAM892). Recipients (SAM599) contained a spectinomycin-resistance cassette (*spc*) with a promoter in the wrong orientation {(P(off)} between two *lox* sites. Transfer and expression of *cre* should cause inversion of the promoter-containing fragment such that the promoter would be driving expression of *spc*{P(on)} (**S1 Fig**). This Cre-mediated recombination is easily detectable by qPCR (**S1 Fig;** Methods) and does not require that the transconjugants be able to form colonies, although viable transconjugants should be spectinomycin resistant.

Cre-mediated recombination enabled us to compare the frequency with which recipients acquired ICE*Bs1* Δ*xis* to that for ICE*Bs1 xis*^+^. We mated ICE*Bs1* Δ*xis cre* (SAM892) with recipients that contained the Cre reporter (SAM599). The acquisition efficiency as measured by CFUs was ~0.01% transconjugants per recipient after two hours of mating (**S2 Fig**). In contrast, the acquisition efficiency as measured by Cre-mediated recombination (qPCR, Methods) was ~5% per recipient after two hours of mating (**Fig 2C**), indicating that using Cre-mediated recombination, we detected 500-fold more inversions than viable transconjugants (**Fig 2D**).

We found that the acquisition efficiency of wild-type ICE*Bs1 cre* (*xis*^+^) (SAM830) as measured by Cre-mediated recombination was ~6% transconjugants per recipient after two hours of mating (**Fig 2C**). This is virtually indistinguishable from the results with ICE*Bs1* Δ*xis* (~5% transconjugants per recipient by the Cre assay). Based on this comparison, we conclude that ICE*Bs1* Δ*xis* actually transferred and integrated into recipients, but those transconjugants were largely inviable and were unable to form colonies.

We found that Cre-mediated inversion underestimated the conjugation efficiency, at least for wild-type ICE*Bs1*. In parallel experiments, the acquisition efficiency of ICE*Bs1* was ~20% transconjugants per recipient after two hours of mating, as measured by CFUs (**S2 Fig**) compared to ~6% as measured by Cre-mediated recombination (**Fig 2C**). After three hours of mating, ~82% of the transconjugants (**Fig 2D**) had a detectable inversion based on the Cre reporter. Virtually every viable transconjugant (>99.8%) recovered was resistant to spectinomycin, indicating that Cre catalyzed an inversion in virtually all transconjugants, but that there was a delay between the initial acquisition of ICE*Bs1* (as measured by CFUs) and the Cre-catalyzed recombination (inversion) event. Together, these results indicate that after two hours the Cre-mediated recombination was detecting ~30% of the viable transconjugants. We suspect that the Cre reporter was also underestimating the acquisition efficiency for ICE*Bs1* Δ*xis*, as it was for ICE*Bs1 xis*+. Together, our results demonstrate that ICE*Bs1* Δ*xis* was transferred to recipients, that the element integrated prematurely into the genome of these transconjugants, and that the vast majority of integrants were not viable. We suspect that loss of viability is likely due to DNA damage from the integration of an element undergoing autonomous rolling circle replication (see below).

### Premature integration of ICE*Bs1* in transconjugants causes chromosome abnormalities

To test if integration of ICE*Bs1* Δ*xis* in naïve hosts caused effects on the host chromosome, we monitored chromosome segregation in transconjugants that had received ICE*Bs1* Δ*xis*. We mated donors and recipients expressing different fluorescent proteins as described above (mApple in recipients; mNeongreen in donors and transconjugants) and then placed mating mixtures on agarose pads containing DAPI to visualize DNA (Methods). We found that ~60% of transconjugants that received ICE*Bs1* Δ*xis* had abnormal chromosome segregation, defined as any division event that did not result in symmetrical segregation of the chromosome to daughter cells, or the appearance of either compacted or elongated nucleoids. In contrast to the ~60% for ICE*Bs1 Δxis*, approximately 20% of ICE*Bs1 xis*+ transconjugants had abnormal chromosome segregation (**Fig 2E-F**). These results indicate that premature integration of ICE*Bs1* in transconjugants caused an increase in chromosome abnormalities.

Detailed characterization of events in transconjugants is technically challenging due to the limited conjugation frequency and the need to distinguish transconjugants from the other two cell types (donors and recipients) present in the mating mix. Below, we describe experiments that overcome these limitations and allow for more detailed characterization of cells in which ICE*Bs1* is integrated and its genes are expressed.

### Use of an ICE*Bs1* Δ*xis* mutant host as a proxy for premature integration in transconjugants

Previous work indicated that activation of ICE*Bs1* that was unable to excise, due to integration in secondary sites, caused a host SOS response and cell death [23], similar to the phenotypes caused by premature integration of ICE*Bs1* Δ*xis* in transconjugants. We used a strain (SAM249) containing ICE*Bs1* Δ*xis* at its normal integration site (*attB*) as a proxy for premature integration of ICE*Bs1* Δ*xis* in transconjugants. This is similar to the donor strain used in conjugation experiments described above, except without providing a functional *xis in trans* and with ICE*Bs1* Δ*xis* integrated at its normal location in the chromosome (*attB* in *trnS-leu2*) Without any *xis* in the cell, activation of ICE*Bs1* Δ*xis* (caused by overexpression of *rapI*) results in de-repression of ICE*Bs1* gene expression and failure to excise from the chromosome, a situation similar to that caused by premature integration in the ICE*Bs1* Δ*xis* transconjugants.

We found that one hour after activation (addition of xylose to induce P*xyl-rapI*) of ICE*Bs1* Δ*xis* in cells growing in defined minimal medium (Methods), cell viability was ~2% of that before activation (**Fig 3**). In contrast, the number of viable cells of ICE*Bs1 xis*+ increased 2-fold under the same conditions.

**Fig 3.**
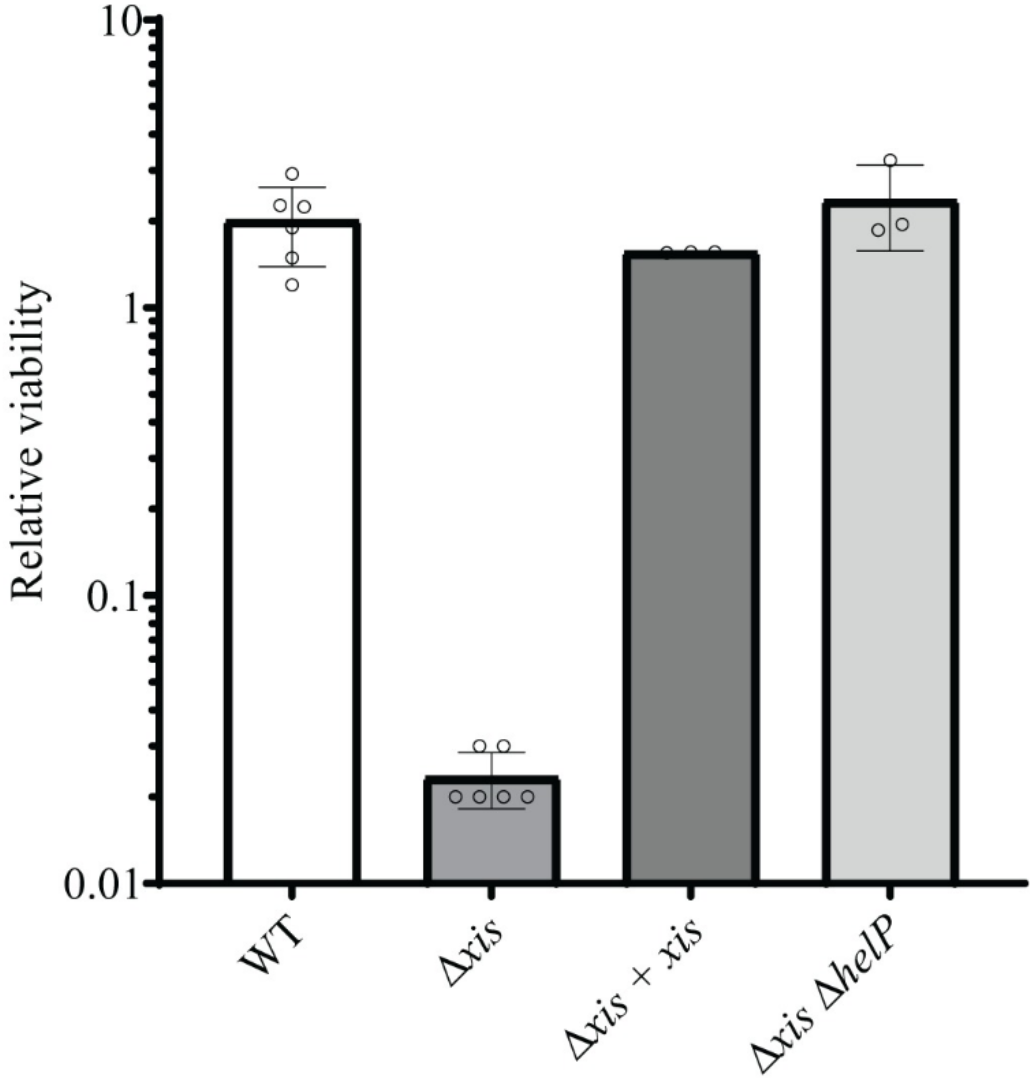
Activation of ICE*Bs1* that is unable to excise from the chromosome causes loss of cell viability. Cells were grown at 37°C in a defined minimal medium containing L-arabinose as a carbon source. Expression of *rapI* (P*xyl-rapI*) was induced by the addition of D-xylose for 60 mins. Viability was determined by CFUs and normalized to the number of CFUs at the time of induction (addition of xylose). ICE*Bs1* WT = AB77; ICE*Bs1* Δ*xis* = SAM249; ICE*Bs1* Δ*xis* + *xis* (i.e., complementation of Δ*xis*) = SAM388; ICE*Bs1* Δ*xis* Δ*helP* (unable to undergo autonomous rolling circle replication) = SAM393. Results are from at least 3 biological replicates. Error bars indicate standard deviation.

Activation of ICE*Bs1* Δ*xis* caused cell death, apparently similar to that observed in ICE*Bs1* Δ*xis* transconjugants. ICE*Bs1* Δ*xis*-containing cells were grown and treated as above and then placed cells on an agarose pad to track the fate of individual cells by time-lapse microscopy (S2 and S3 Movie). We found that during a 3 hour period following 1 hour of growth in ICE-inducing conditions, cell lysis occurred in ~70% of cells containing ICE*Bs1* Δ*xis* compared to ~ 5% of those containing ICE*Bs1 xis*+.

We found that the loss of viability of cells with ICE*Bs1* Δ*xis* was due to loss of *xis* and not some unexpected effect on downstream genes or a *cis*-acting element in ICE*Bs1*. We provided *xis in trans* from an ectopic locus (SAM388), thereby restoring excision. Viability was also restored to levels seen for cells with ICE*Bs1 xis+* (**Fig 3**), analogous to the restoration of the timing of integration observed when ICE*Bs1* Δ*xis* transfers into recipients expressing *xis* (**Table 1, Fig 2A**). Together, these results indicate that ICE*Bs1* Δ*xis* is a suitable proxy for premature integration of ICE*Bs1* Δ*xis* in transconjugants, that loss of cell viability is due to the inability of the element to excise from the chromosome, and that the terminal phenotype is cell lysis.

### Cell death precedes lysis

Although cell lysis was the terminal phenotype, we found that cell death preceded lysis. We used staining with propidium iodide (PI) as an indicator of cell death (Methods; [24]) and tracked the ability of cells to take up PI and cell lysis using microscopy. We found that at the time ICE*Bs1* Δ*xis* cells (SAM427) became PI-positive there was an appreciable decrease (~25%, n = 20) in cell surface area (**S3A and S3B Fig)**, consistent with cell membrane permeability and loss of the proton motive force [25]. Cell lysis occurred on average by 12 minutes after staining with PI and the change in surface area. Together these results indicate that cell death caused by activation of ICE*Bs1* Δ*xis* occurred before cell lysis. We suspect that loss of the proton motive force likely activates *B. subtilis* autolysins which then cause cell lysis [26,27].

The inability to excise does not immediately explain the loss of cell viability in ICE*Bs1* Δ*xis* and indicates that there is another function of ICE*Bs1* biology that is lethal when the element is activated and unable to excise, or when an active element is integrated prematurely in transconjugants.

### Activation of ICE*Bs1* Δ*xis* is lethal due to rolling circle replication of the element in the host chromosome

When activated, ICE*Bs1* undergoes unidirectional rolling circle replication that initiates from its origin of transfer (*oriT*) [8,15]. When the element is unable to excise due to deletion of the attachment site at the right end (Δ*attR*), this replication initiates in the integrated ICE and extends into the chromosome [8]. Replication of ICE*Bs1* requires three element-encoded functions: the relaxase NicK [15], the origin of transfer (*oriT*) that is nicked by the relaxase [15] and also functions as an origin of replication [8,28], and the helicase processivity factor HelP [28]. In an ICE*Bs1* Δ*helP* mutant, nicking occurs at *oriT* [15], but the ICE DNA is not unwound and cannot be replicated [28].

We found that the lethality caused by activation of ICE*Bs1* Δ*xis* was likely due to replication of the integrated element. There was no cell death in the ICE*Bs1* Δ*xis* Δ*helP* mutant (SAM393) (**Fig 3**), indicating that replication, or at least DNA unwinding, was causing cell death following activation of the excision-defective element. These results demonstrate that the loss of viability and subsequent lysis (above) caused by an activated ICE*Bs1* Δ*xis* was most likely due to rolling circle replication of an element that is unable to excise. Further, cell lysis was the terminal phenotype, consistent with results above for early integration of ICE*Bs1* Δ*xis* in transconjugants.

### Activation of ICE*Bs1* Δ*xis* induces the SOS response

Rolling circle replication of integrated ICE*Bs1* originates from *oriT*, extends into the chromosome, and generates ssDNA [15,28]. Iterative replication, that is multiple initiation events at the newly synthesized *oriT*, could also lead to a fragile region of the chromosome and potential arrest of chromosome replication forks that had initiated at the chromosomal origin, *oriC*. Both the increase in ssDNA and arrest or collapse of chromosomal replication forks could trigger the RecA-dependent SOS response. RecA is loaded onto ssDNA, displacing the host single-strand binding protein SSB, and the RecA-ssDNA nucleoprotein filament stimulates autocleavage of LexA, resulting in de-repression of genes in the LexA regulon (SOS response).

We monitored induction of the SOS response in single cells using a transcriptional fusion of the gene encoding the fluorescent protein mNeongreen to the promoter for *yneA* (*yneA-mNeongreen*), a gene that is repressed by LexA and activated during the SOS response [29]. We found that the SOS response was activated two hours after induction of the excision-defective ICE*Bs1* Δ*xis*. The median fluorescence intensity of the population increased ~20-fold compared to the uninduced ICE*Bs1* Δ*xis*, with ~83% of the population responsible for fluorescence (**Fig 4A-B**). In contrast, we found that two hours after activation of the excision competent ICE*Bs1* (*xis*+), the median fluorescence intensity of the population increased ~2-fold compared to the uninduced ICE*Bs1 xis+*, with ~10% of the population responsible for fluorescence (**Fig 4A-B**). These results indicate that activation of ICE*Bs1* Δ*xis* causes activation of the SOS response in the majority of cells in the population and are consistent with previous findings that ICE*Bs1* that is unable to excise from the chromosome causes induction of the SOS response [23].

**Fig 4.**
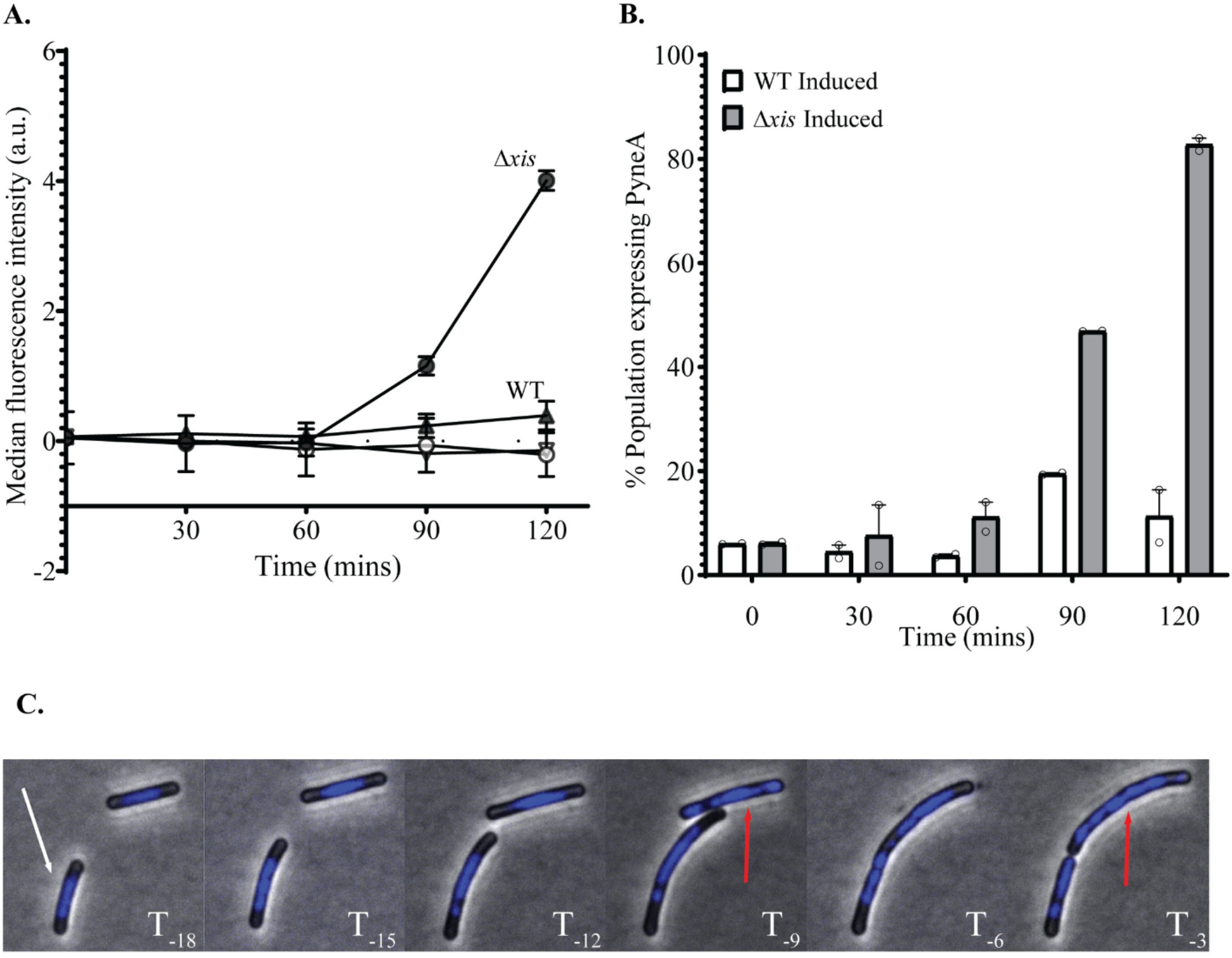
Activation of ICE*Bs1* that is unable to excise causes DNA damage. **A)** Cells contained the SOS reporter *PyneA-mNeongreen*. Strains were grown at 37°C in a defined minimal medium with L-arabinose as the carbon source. Expression of *rapI* (P*xyl-rapI*) was induced by the addition of D-xylose. Samples were then spotted onto agarose pads and imaged at the indicated times. The average median fluorescent intensity of individual cells from at least 2 experiments is plotted as a function of time after addition of xylose. WT = SAM426; Δ*xis* = SAM427. WT Induced = 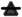; WT Uniduced = 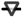; Δ*xis* Induced = 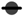; Δ*xis* Uninduced (no xylose) = 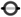. Error bars indicate standard error of the mean. **B)** Fraction of population gated as SOS+ based on ICE*Bs1* uninduced distribution. Same experiment as **A**. Points represent mean from an individual experiment. Error bars indicate standard error of the mean. **C)** Induction of ICE*Bs1* Δ*xis* (SAM427) causes chromosome abnormalities. Experimental conditions were the same as in panel 4**A**. After 60 mins of ICE*Bs1* induction, cells were washed and spotted onto agarose pads that contained CH medium and DAPI. Images were captured every 3 mins. The indicated times (T_-x_) indicate the number of mins before lysis of the cell indicated by the white arrow in the bottom left of the first panel. Red arrows indicate cells with abnormal chromosomes right before lysis. Note: Both cells present lyse at T_-18_.

### Activation of ICE*Bs1* Δ*xis* causes chromosome abnormalities

We found that activation of the excision-defective mutant (ICE*Bs1* Δ*xis*) caused chromosome abnormalities, analogous to those caused by premature integration of ICE*Bs1* Δ*xis* in transconjugants. We grew cells containing ICE*Bs1 xis+* (SAM426) or ICE*Bs1* Δ*xis* (SAM427) in defined minimal medium, activated ICE*Bs1* by overexpression of *rapI* (from P*xyl-rapI*) for one hour, then removed the activator and placed cells on an agarose pad that supported rapid growth (doubling time of ~20 minutes). To observe the nucleoid (chromosomal DNA) and septum formation, agarose pads contained DAPI (DNA staining) and FM4-64 (membrane staining) and cells were visualized every three minutes for three hours (Methods).

We found that the majority (~92%) of cells with an activated ICE*Bs1* Δ*xis* had aberrant nucleoids and-or defects in chromosome segregation (**Table 2; Fig 4C; S4A–S4C and S4G–S4H Fig, S4 Movie**), including approximately 30% of cells with aberrantly elongated nucleoids (**S4D–S4F Fig**). We defined aberrant chromosome segregation as nucleoids that had an asymmetric distribution between daughter cells. Daughter cells were defined as cells that existed for at least one frame after division and prior to lysis.

**Table 2.**
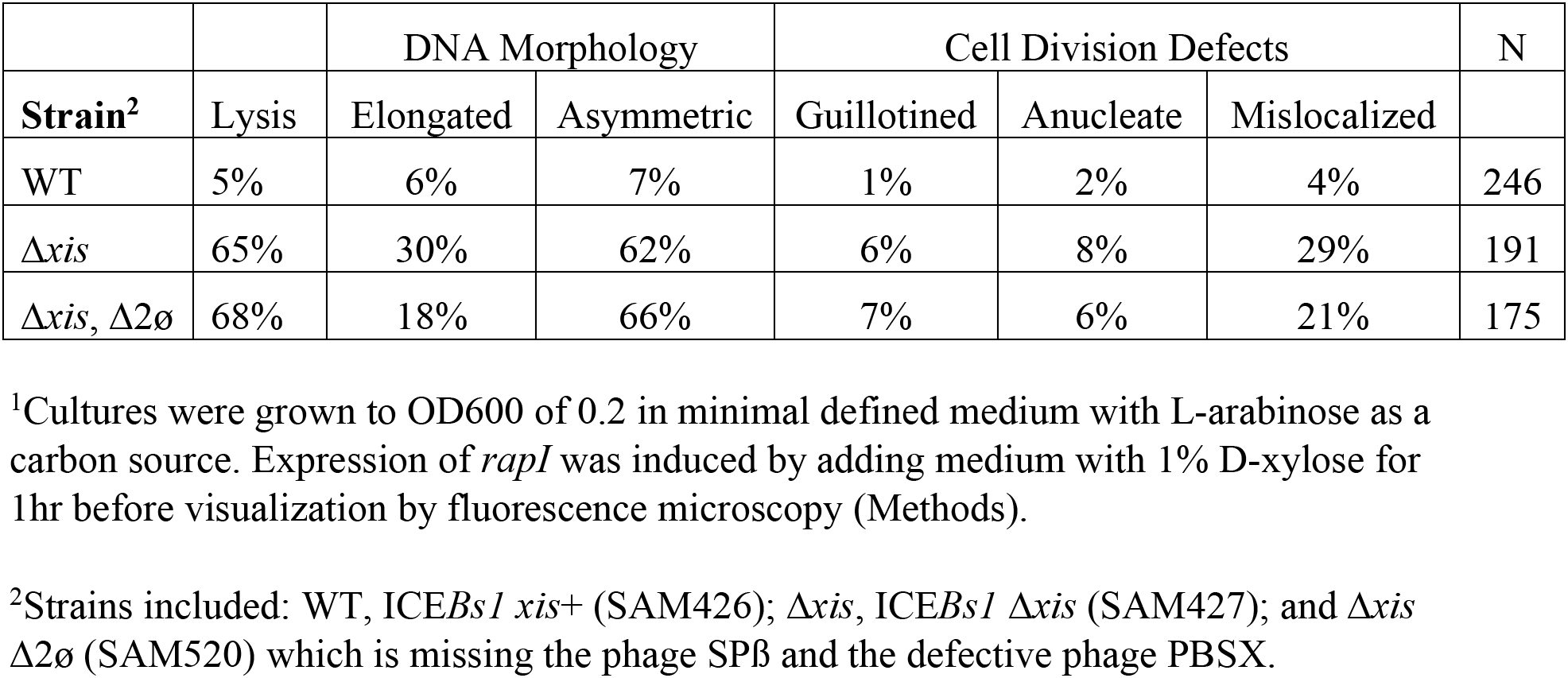
DNA morphology and cell division defects following expression of ICE*Bs1*^1^.

In addition to abnormal chromosome segregation and elongated nucleoids, many cells had defects in cell division, including mislocalized septa (septum formation not at mid-cell), guillotined chromosomes (**S4D–S4F Fig**), and anucleate cells (**S4G–S4H Fig**). We found that 29% of cells with an activated ICE*Bs1* Δ*xis* had mislocalized septa, 6% had guillotined chromosomes, and 8% were anucleate. For those cells that were anucleate, we do not have the temporal resolution to determine whether this was due to defects in chromosome segregation or chromosome degradation. However, given the large fraction of cells that had abnormal septa, we believe most of the anucleate cells could be due to segregation defects. Additionally, approximately 65% of the population lysed during the course of the experiment (**Table 2**).

These effects caused by activation of ICE*Bs1* Δ*xis* were markedly more prevalent than those caused by activation of ICE*Bs1* that was able to excise (ICE*Bs1 xis*+). Of the cells in which ICE*Bs1* was activated and able to excise (SAM426), only ~5% lysed, <15% had aberrant-looking chromosomes, and <10% had any apparent defect in cell division, compared to ~65% lysed, ~92% with chromosome abnormalities, and ~43% with cell division defects in cells with ICE*Bs1* Δ*xis* (**Table 2**). Together, these results indicate that activation of ICE*Bs1* Δ*xis* (unable to excise from the chromosome) causes severe perturbations in chromosome organization and segregation, cell division, and ultimately causes cell death.

### Chromosome abnormalities and cell lysis caused by activation of ICE*Bs1* Δ*xis* were not due to activation of phage

We found that the chromosome abnormalities and cell lysis caused by activation of ICE*Bs1* Δ*xis* occurred in the absence of the functional phage SPß and the defective phage PBSX that are present in the *B. subtilis* chromosome. Both SPß and PBSX are activated during the SOS response [29] and cause cell lysis [30–33]. In a strain devoid of these two elements (SAM520), activation of ICE*Bs1* Δ*xis* still caused chromosome abnormalities and cell lysis (**Table 2; S5 Movie**). Based on these results, we conclude that activation of ICE*Bs1* Δ*xis* causes chromosome abnormalities and cell lysis independently of these phage or phage-like elements.

There was always a fraction of cells in our experiments that had no detectable chromosome abnormalities, appeared to grow normally, and in contrast to most other cells in the population, did not lyse. This apparent normalcy could be because these cells were resistant to the effects of ICE*Bs1* Δ*xis*, ICE*Bs1* Δ*xis* was not activated, or the element had been lost. We used a fluorescent reporter fused to the major ICE*Bs1* promoter (P*xis*-*mApple*; at an ectopic location in the chromosome; i.e., not in ICE*Bs1*) that allowed us to distinguish between these possibilities. There is little to no fluorescence from P*xis-mApple* in cells with ICE*Bs1* that is integrated in the chromosome and fully repressed (i.e., not activated) due to repression by the element-encoded repressor ImmR. Expression from P*xis* is high in cells without ICE*Bs1* and intermediate in cells with ICE*Bs1* that had been activated (de-repressed) **(S5 Fig)**.

We found that virtually all cells that had normal-looking chromosomes and that did not lyse had little or no detectable expression of P*xis-mApple* (**S6 Movie, S6 Fig**), indicating that ImmR was present and fully active. Based on these results, we conclude that these cells still contained ICE*Bs1* Δ*xis*, but that the element had not been activated. There were some cells that had activated ICE*Bs1* and had chromosome abnormalities but did not lyse (**S6 Fig**). It is possible that these cells were destined to lyse, but not during the three hours of our observations.

## Discussion

We used a combination of population and single-cell analyses to study integration of ICE*Bs1* in transconjugants. We found that integration occurs several generations after initial transfer, early integration is lethal to most cells, and lethality is likely due to rolling circle replication from ICE*Bs1* that had integrated into the chromosome. Furthermore, based on our results, we conclude that premature integration of ICE*Bs1* in transconjugants causes phenotypes similar to those caused by activating an element that is unable to excise from the chromosome. In both cases, rolling circle replication from ICE*Bs1* in the chromosome causes a host SOS response, chromosome abnormalities, and cell death.

Our results demonstrate that robust viability and fitness of transconjugants depends on repression of replication before or concomitant with integration. If integration happens while rolling circle replication is ongoing, there are chromosome abnormalities and death of transconjugants. We conclude that ICE*Bs1* normally links integration, which is required for stable acquisition and maintenance of an element in a population [19], to the cessation of autonomous ICE*Bs1* replication. The ability of ICE*Bs1* to remain extrachromosomal and replicate autonomously is beneficial for the establishment and spread of the element, and for the hosts if the presence of the element confers a fitness advantage. Replication of ICE*Bs1* upon transfer to a new host allows for the element to segregate during growth and division of the transconjugants and for the transconjugants to become donors as soon as the ICE genes needed for conjugation are expressed. This is especially critical for transfer of the element in chains of cells and cells growing in communities [20,34–36]. Below, we discuss the linkage between integration and cessation of autonomous replication and postulate that it is a conserved regulatory feature of integrative mobile genetic elements, including ICEs and temperate phages that integrate into the host genome.

### Excisionases, required in donors, are also important in transconjugants

There is a large body of work demonstrating the importance of excisionases (recombination directionality factors) for the excision of ICEs and prophages from the chromosome of host cells [37]. However, it is largely assumed that excisionases are not important for establishment of an element after it enters a naïve host. We found that in addition to its well-known and expected role in excision from the chromosome [19], the ICE*Bs1* excisionase is critical following conjugative transfer of the element to a new host (transconjugant) and that it acts to affect the timing of stable integration.

Upon entry into a new host, most ICE*Bs1* genes are expressed as there is no repressor (ImmR) present and the major promoter P*xis* is on (**Fig 1**). This prevents integration and enables new transconjugants to serve as donors while the element is still extrachromosomal and expressing genes needed for autonomous rolling circle replication and conjugation, thereby increasing transfer of the element in a population [20,34–36]. This de-repressed gene expression upon transfer to a naïve host is analogous to zygotic induction of phage lambda following its transfer to a new, non-lysogenic host via conjugation [38].

After transfer to a new host, ICE*Bs1* genes encoding the repressor (*immR*), anti-repressor (*immA*), and integrase (*int*) are also expressed. Once enough ImmR accumulates, it activates its own promoter and represses transcription from P*xis* [18,19]. This regulatory circuit also stimulates transcription of *int*. Based on the mechanisms of action of integrases and excisionases and comparison to analogous systems, integration of ICE*Bs1* would occur following accumulation of the integrase and depletion of the excisionase. Depletion of the previously made excisionase occurs by dilution during cell growth and degradation if the protein is unstable. Although we have not yet tested it directly, we suspect that Xis is degraded by host proteases, thereby enabling integration shortly after repression of P*xis* by ImmR, analogous to degradation of λ Xis [39].

Once repression of P*xis* occurs, the ICE*Bs1* genes required for rolling circle replication are no longer expressed and autonomous replication will cease. In this way, integration of ICE*Bs1* into the chromosome of its new host is enabled, either concomitant with or shortly after the cessation of replication. This coordinated timing affects the fitness of ICE*Bs1* and host cells in several ways: 1) It allows replication and maintenance of ICE*Bs1* in the growing transconjugants prior to integration; 2) It allows early transconjugants to become donors before integration; and 3) It ensures that when the element does integrate into the genome, most of the integrants will be stable and viable with no autonomous replication of the integrated element.

### Regulation and timing of integration of other ICEs

It is not known if premature integration of other ICEs causes phenotypes similar to those described here. However, there are several ICEs that are known to undergo autonomous replication, including *Tn916* (*Enterococcus faecalis*), ICE*clc* (*Pseudomonas knackmussii B13*), ICE*Hin*1056 (*Haemophilus influenzae*), ICE*MlSym^R7^*(*Mesorhizobium loti^R7A^*), ICE*SXT/R391* (*Vibro cholera/Providencia rettgeri*), Tn*GBS1/TnGBS2* (*Streptococcus agalactiae*), and ICE*St*1/ICE*St*3 (*Streptococcus thermophilus*). If autonomous replication of any of these ICEs occurs in transconjugants, then it seems likely that replication ceases prior to or concomitant with integration. We suspect that this is also true in the initial host if the element were to excise and then reintegrate. Additionally, some of these, including ICE*Hin*1056 and *TnGBS1/TnGBS2*, are known to exist transiently as replicating elements in transconjugants, at least for a few generations prior to integration. These properties indicate that integration likely occurs after replication is halted.

#### ICE*Hin*1056

Although it was not known to be an ICE at the time, prior work characterizing what is now known as ICE*Hin1056* revealed that donors, without detectable free plasmid during growth, were able to transfer antibiotic resistance to recipients. The resulting transconjugants transiently contained plasmid DNA and were capable of efficient transfer [40,41]. These initial studies indicated that at least some ICEs likely exist as extrachromosomal plasmids for several generations before integrating into the nascent host chromosome and that while plasmids, they were capable of efficient transfer to other cells.

#### Tn*GBs1*/Tn*GBS2*

ICEs in the *TnGBS* family differ from most characterized ICEs in that they use a DDE transposase rather than a phage-like integrase. The limited host range of Tn*GBS* ICEs was found to be due to the inability of the element to replicate in species other than *S. agalactiae*. Further, qPCR analysis of the ratio of integrated ICE to circular ICE revealed that Tn*GBS* ICEs remained in the circular form in transconjugants for some time before integration into the chromosome, and that the timing of integration varied between elements [14].

#### Other ICEs

For some ICEs, for example, ICE*SXT*, ICE*MlSym^R7A^*, R391 and ICE*Bs1*,autonomous replication is critical for the maintenance of the element in donors following activation and excision [8,13,16]. In addition, for ICE*Bs1* and ICE*MlSym^R7A^*, replication is important in the generation of stable transconjugants [16,28]. Autonomous replication of ICE*clc* in donors increases the conjugation efficiency [7,10], and the element remains extrachromosomal in transconjugants for up to 10 hours after transfer [7]. For several ICEs, it has been shown that dysregulation between replication and integration results in instability of the element [8,9,13,16,19]. Together, the knowledge that various ICEs replicate autonomously indicates that proper regulation of integration is likely important for the stability of the element and survival of host cells.

### Mechanisms for coupling integration with cessation of autonomous replication

There are at least three strategies used by ICEs to couple integration with cessation of autonomous replication: 1) Coordinated repression of excision and replication functions; 2) Excisionase-mediated activation of genes needed for replication; and 3) Physical separation of replication genes from their promoter upon integration.

#### 1) Coordinated repression of excision and replication functions

ICE*Bs1* regulatory hierarchy involves the co-transcription of genes needed for excision, replication, and conjugation [8,17–19,28]. The gene for the excisionase is upstream of those for replication and conjugation. As of yet, there are no published data on possible differential regulation of expression of excision and replication genes for ICE*Bs1*. However, we note that a review of a transcriptomics study of *B. subtilis* [42] indicates an additional promoter downstream of the excisionase, driving expression of genes needed for replication and conjugation. The presence of this promoter could provide additional regulatory inputs that might affect the timing of integration and cessation of replication. Additionally, we suspect that the ICE*Bs1* excisionase (and probably others) is unstable, providing tighter temporal linkage between the repression of *xis* expression, disappearance of the protein, and the ability of the element to stably integrate.

#### 2) Excisionase-mediated activation of genes needed for replication

ICE*MlSym^R7A^* appears to use a similar regulatory hierarchy as ICE*Bs1*. Variants of ICE*MlSym^R7A^* are found in other *Mesorhizobium* species and have a similar regulatory hierarchy [43,44]. Additionally, for these ICEs the excisionase has been found to act as a transcription factor, activating the expression of downstream genes required for the transfer of the elements. This indicates that ICEs of the *Mesorhizobium* spp., may use a similar method of regulation observed for CTn*DOT* of *Bacteroides thetaiotamicron*, in which the excisionases moonlight as transcription factors, activating the expression of genes needed for conjugative replication, thereby directly linking Xis to the regulation of replication [44,45].

#### 3) Physical separation of replication genes from their promoter upon integration

*Tn916* of *Enterococcus faecalis* employs a regulatory hierarchy that involves physical separation of promoters from genes needed for replication and conjugation [46]. Excision of the element orients the promoter such that transfer and replication genes are transcribed. Conversely, integration separates the genes needed for replication and conjugation from the promoter(s) that drive their expression, thereby preventing further autonomous replication of the element.

## Conclusions

This linkage between integration and the absence of autonomous replication likely pertains to ICEs that undergo autonomous replication and many temperate phages. Temperate phages that infect naïve hosts and integrate into the genome to form lysogens typically do so prior to the start of autonomous replication. This is typically achieved by a hierarchy of phage genes expressed at different times. Phage genes required for replication are typically expressed after the early genes and only if the phage enters the lytic cycle [47–50].

In summary, our work provides a new perspective on ICE regulation as it pertains to transconjugants. We uncovered a critical role for the excisionase in transconjugants and revealed the importance of regulatory mechanisms that delay integration until the cessation of replication. Because of the deleterious effects of integration before the cessation of replication, it is likely that most, and perhaps all integrative elements that undergo autonomous replication have a mechanism that enables integration only after replication is no longer possible. Because of the prevalence of ICEs, their powerful ability to transfer genes, and their recent uses in genetic analyses and engineering [51,52], it is important to understand ICE regulatory mechanisms and events in new hosts. This understanding could inform strategies to reduce the spread of undesirable elements (e.g, encoding clinically relevant antibiotic resistances) and facilitate the use of ICEs for engineering a wide range of microbes.

## Methods and Materials

### Media and bacterial growth

Cells were grown in LB medium or defined S7_50_ minimal medium with 0.1% glutamate [53] and a carbon source, typically 1% L-arabinose, as indicated. Serial dilutions and resuspensions for plating viable CFUs were done in 1x Spizizen salts [54]. For experiments, indicated strains were colony purified from frozen (−80°C) glycerol stocks on LB plates with the appropriate antibiotics. A single colony was picked and used to start a culture which was grown to mid-exponential phase. An aliquot was then diluted into fresh medium and allowed to grow to an appropriate culture density in mid-exponential phase for each experiment. Cells were typically grown in flasks at 37°C with shaking (250 RPM).

Where indicated, ICE*Bs1* was induced by the addition of D-xylose (1%) to cause overexpression of *rapI* from a xylose-inducible promoter in strains containing either *amyE*::{(P*xyl-rapI*) *spc*} [55] or *lacA*::{(P*xyl-rapI*), *tet*} [8]. Where indicated, cells were also grown with L-arabinose to induce expression of the xylose transporter. Expression from the LacI-repressible-IPTG-inducible promoter *Pspank* was induced with 2 μM IPTG.

Antibiotics used included: kanamycin (5 μg/ml); chloramphenicol (5 μg/ml); tetracycline (12.5 μg/ml); spectinomycin (100 μg/ml); streptomycin (100 μg/ml); and 0.5 μg/ml erthyromycin plus 12.5 μg/ml lincomycin to select for macrolide-lincosamide-streptogramin B (MLS) resistance conferred by *mls* (*ermB*).

### Bacterial strains and alleles

All *B. subtilis* strains (**Table 3**) are derivatives of JH642 (AG174; [56,57]and contain *trpC* and *pheA* mutations (not indicated in the table). Most strains were constructed by natural transformation to introduce appropriate alleles. In some cases, as indicated, ICE*Bs1* was transferred into strains by conjugation. ICE*Bs1* in most strains contained a gene conferring resistance to an antibiotic, often *kan* (kanamycin resistance), inserted in the *rapI-phrI* locus {e.g., Δ(*rapI-phrI*)*342*::*kan*} [17]. Strains cured of ICE*Bs1* [17] are indicated as ICE*Bs1*^0^. Almost all alleles were first constructed in so-called ‘clean’ strains that were typically wild-type *B. subtilis* with few if any other alleles. These were verified by phenotype, diagnostic PCR, and sometime sequencing. Alleles were then transferred to other strains to build strains with complex genotypes.

**Table 3.**
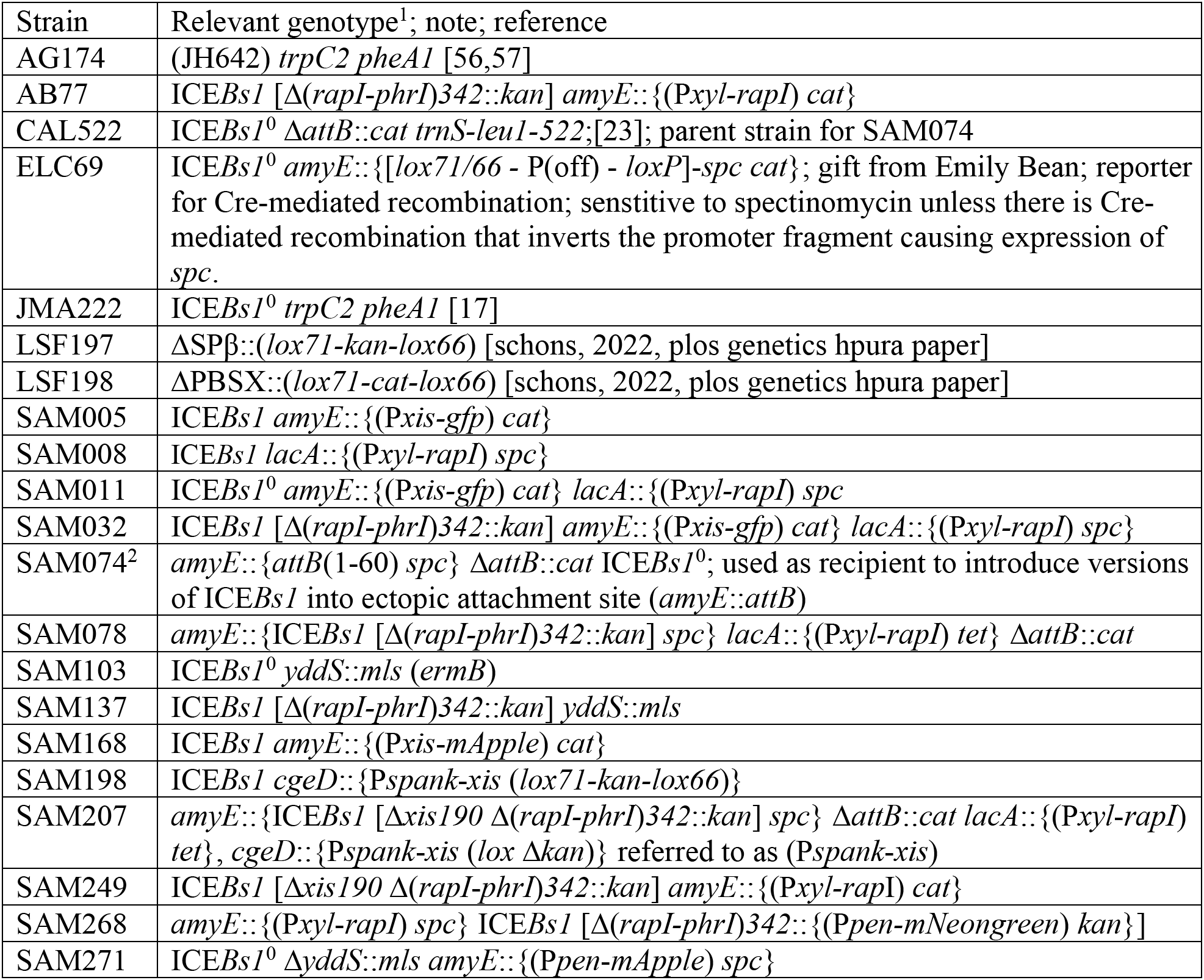
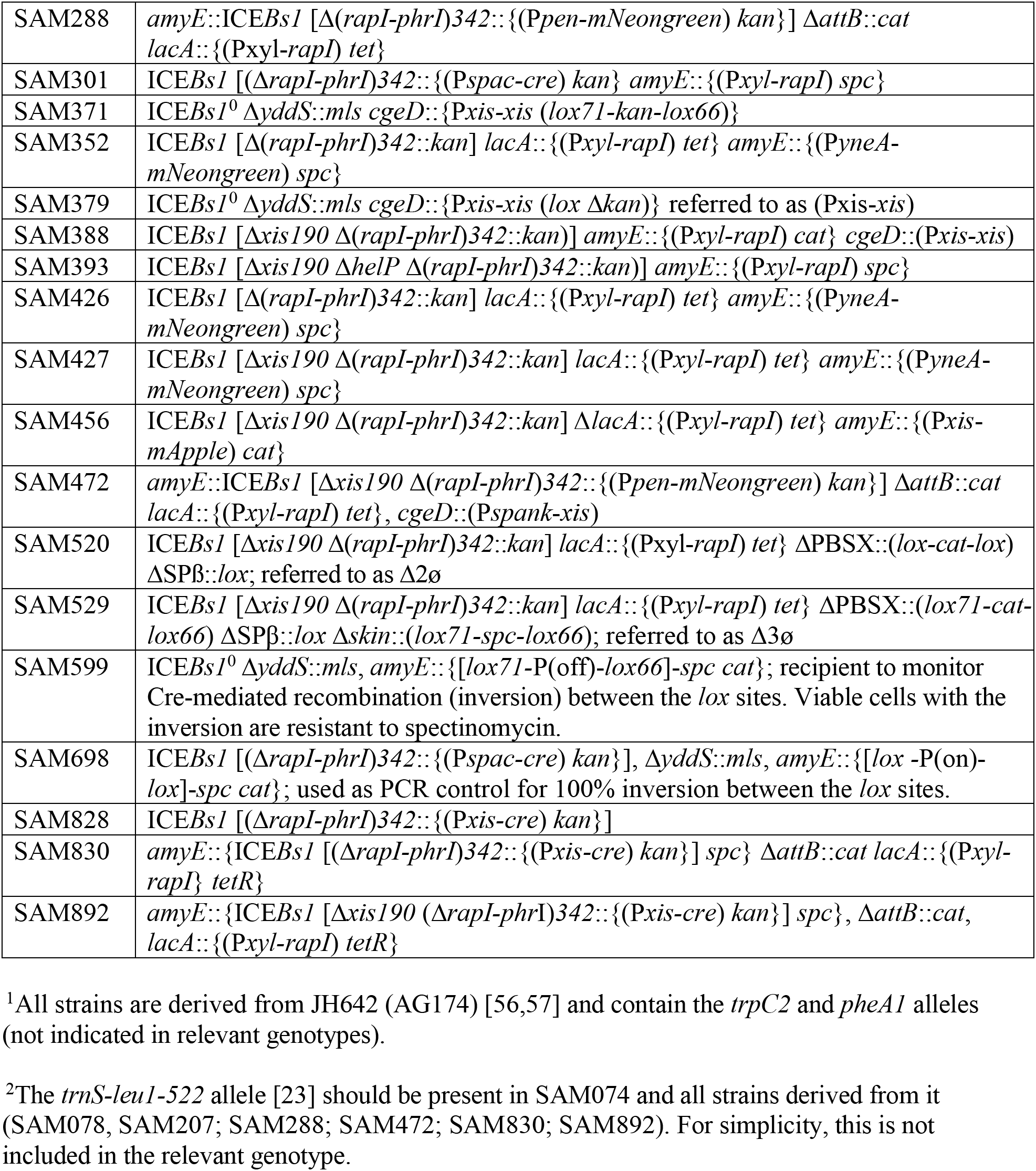
*B. subtilis* strains used.

#### amyE::attB

SAM074 contains the chromosomal attachment site for ICE*Bs1 (attB*) cloned into the nonessential gene *amyE* (*amyE*::*attB*) It is cured of ICE*Bs1*, and contains a deletion of *attB* from its normal location (Δ*attB*::*cat*) that was described previously [23]. *amyE*::(*attB spc*) was made by cloning *attB* into the *amyE* integration vector pDG364 [58]Oligonucleotides that contained the 60-bp *attB* were annealed and 25-bp of random sequence on the flanking ends were added and cloned into pDG364. The *cat* gene in pDG364 was then replaced with *spc* (spectinomycin resistance), that had been amplified from pMMB856 [55]. This plasmid (pSAM071) was then used to introduce *attB* into the *B. subtilis* chromosome at *amyE* by transformation into strain CAL522 (Δ*attB*::*cat*), selecting for resistance to spectinomycin.

Various versions of ICE*Bs1* were introduced into SAM074 (*amyE*::*attB*, Δ*attB::cat*) by conjugation with the relevant donor and SAM074 as recipient and selecting for transconjugants. Strains were verified by PCR and sequencing of relevant alleles. Strains constructed in this way include: SAM078; SAM207; SAM288; SAM472; SAM830; and SAM892.

#### yddS::mls

*yddS* is ~5 kb from *attB* and *yddS::mls* was used to distinguish recipients and transconjugants from donors, with *mls* serving as a counter-selective marker (MLS resistance) to prevent growth of donors, and a sequence for PCR that was present in only recipients and transconjugants. The allele contains an insertion of *mls* at base pair 22 of the 1,311 bp *yddS* open reading frame and was made by PCR amplifying ~2 kb of DNA from upstream and downstream of the insertion site and assembling these (Gibson isothermal assembly [59]) such that they flanked a DNA fragment containing *mls* that had been amplified from pCAL215 [18]. This fragment was then cloned between the EcoR1 and KpnI sites in pUC19, generating pSAM095, which was then used to transform *B. subtilis* selecting for MLS resistance.

#### Ectopic expression of *xis*

Constructs for ectopic expression of ICE*Bs1 xis* included *cgeD::(Pspank-xis*) and *cgeD*::(P*xis*-*xis*). Both were made with the plasmid pSAM178, a vector for integrating cloned DNA at *cgeD*. pSAM178 contains the *gatB/yerO* terminator adjacent to a multi-cloning site, *kan* (flanked by *lox71* and *lox66* sites for removal of *kan* by Cre-mediated recombination [60,61] as indicated, between sequences from *cgeD* in the pMMB124 [18] backbone. Plasmid pSAM189 contains *Pspank-xis* [19] in the pSAM178 backbone. pSAM276 contains P*xis-xis* amplified from its endogenous location from 158 bp upstream of the *xis* open reading frame to 33 bp into *ydzL*, the gene downstream from *xis*, in the pSAM178 backbone. Both constructs were introduced into *B. subtilis* by natural transformation and selection for kanamycin resistance. Strains were confirmed by diagnostic PCR and checking for chloramphenicol sensitivity (loss of the plasmid backbone). *kan* was then deleted from these strains by introducing *cre* on a temperature sensitive plasmid that confers MLS resistance and expresses the Cre recombinase (pSAM097). Strains were then cured of pSAM097 by switching to growth at a non-permissive temperature without selection. Strains with ectopic expression of *xis* include: <u>SAM198, SAM207, SAM371, SAM379, SAM388, and SAM472</u>.

#### Fusions to *mNeongreen, gfp, and mApple*

*amyE*::P*yneA-mNeongreen* (in strains SAM426, SAM427) was used to monitor SOS induction. The fusion contains sequences extending from the first nucleotide to 151 bp upstream of the *yneA* open reading frame, including the promoter region [62,63]. This was fused to *mNeongreen* and flanked by sequences for recombination into *amyE*, including *spc*, by linear Gibson isothermal assembly [59]. The resulting construct was then transformed into *B*. subtilis selecting for spectinomycin resistance to generate strain SAM352. Genomic DNA from SAM352 was used to move *amyE*::{(*PyneA-mNeongreen*) *spc*} into various strains by transformation selecting for spectinomycin resistance.

P*pen-mNeongreen* in ICE*Bs1* (ICE*Bs1* Δ(*rapI-phrI*)*342*::{(P*pen-mNeongreern*) *kan*} in SAM268; SAM288; SAM472) was used to visualize ICE*Bs1* in single cells. The constitutive promoter P*pen* was amplified from pMMB1010 [20] (with a *spoVG* ribosome binding site in one of the PCR primers downstream of P*pen* for efficient translation of *mNeongreen*) and fused to *mNeongreen*. This fusion was flanked by sequences upstream and downstream from the endpoints of Δ(*rapI-phrI*)*342* and included *kan* [17] using Gibson isothermal assembly. This was then transformed into *B. subtilis* (ICE*Bs1*+), selecting for kanamycin resistance and verifying that the reporter was in ICE*Bs1*.

*amyE*::P*pen-mApple* (SAM271) was used to distinguish recipients and transconjugants from donors in conjugation experiments visualizing single cells. The constitutive promoter P*pen* was amplified from pMMB1010 [20], with the ribosome binding site from *spoVG* (as above), and placed upstream of *mApple*. This fragment was then cloned into the *amyE* integration vector pDR160 [21], replacing the xylose-inducible cassette, followed by integration of P*pen-mApple* at *amyE* by double-crossover homologous recombination (between the *amyE* fragments in pDR160) selecting for spectinomycin resistance.

*amyE*::P*xis-gfp* (in SAM005, SAM011, and SAM032) and *amyE*::P*xis*-*mApple* (in SAM168, SAM456) were used to report on expression levels of P*xis*, as a reflection of the amount of active ImmR (the ICE*Bs1* repressor) in the cells. P*xis* was amplified from its endogenous location from 215 bp to 4 bp upstream of the *xis* open reading frame, fused to *gfp* or *mApple*, and cloned into the *amyE*-integration vector pDG364[58], generating plasmids pSAM003 (P*xis-gfp*) and pSAM159 (P*xis-mApple*). These plasmids were transformed into AG174 or JMA222 by natural transformation selecting for chloramphenicol resistance generating strains SAM005 (P*xis-gfp*) and SAM168 (P*xis*-*mApple*). Strains were confirmed by diagnostic PCR and sequencing. Genomic DNA was used to move alleles by transformation to build other strains, including SAM011 and SAM456.

#### ΔSPß::(*lox-kan-lox*), ΔPBSX::(*lox-cat-lox*), Δ*skin*::(*lox*-*spc-lox*) in strains SAM520 and SAM529

Deletion-insertion mutations of the prophage SPß and the defective phages PBSX were described [64]. These essentially delete the indicated phage and insert an antibiotic resistance gene flanked by *lox* sites. The Δ*skin*::(*lox-spc-lox*) mutation was constructed similarly, by PCR amplifying genomic DNA from upstream and downstream of *skin* and then using Gibson assembly to place these PCR products fragments on either side of *lox-spc-lox*. This was then transformed into *B. subtilis*, selecting for spectinomycin resistance, resulting in deletion of *skin* and insertion of *lox-spc-lox*. Genomic DNA was used to move alleles by transformation to build strains SAM520 and SAM529. Where indicated, the antibiotic resistance gene was deleted by Cre-mediated recombination as above.

#### ICE*Bs1* Δ(*rapI-phrI*)*342*::P*xis-cre kan* in strains SAM828, SAM830, and SAM892

*cre*, encoding the Cre recombinase, was cloned into ICE*Bs1* and used to monitor transfer of the element to cells that might otherwise not remain viable. Transconjugants could be detected with a reporter that would recombine in the presence of Cre. To construct P*xis-cre* in ICE*Bs1*, the promoter P*xis* was amplified from its endogenous location from 158 bp upstream of and to 4 bp into the *xis* open reading frame. *cre* was amplified from pDR244 [65] and the ribosome binding site from *spoVG* [66] was contained on one of the primers and placed upstream of *cre* to increase translation efficiency. Two DNA fragments, one upstream and one downstream of the endpoints of Δ(*rapI-phrI*)*342*, were amplified, including *kan*, and the four fragments were assembled using linear Gibson isothermal assembly. This was then introduced into ICE*Bs1* by transformation selecting for kanamycin resistance.

The reporter in recipient cells (SAM599) that was used to indicate acquisition of *cre* and subsequent Cre recombinase activity (S1 Fig) was constructed by Emily Bean (original strain ELC69). It consists of a promoter flanked by *lox* sites [60] such that recombination would result in a stable inversion [61]. The *lox71*-promoter-*lox66* fragment was upstream from *spc*. The initial orientation is such that the promoter is pointed away from *spc* {P(off)}. Cre-mediated recombination would invert the promoter-containing fragment such that the promoter would be driving expression of *spc* {P(on)}. Viable transformants that underwent Cre-mediated recombination would be spectinomycin resistant. Recombinants are also easily detectable by PCR with primers diagnostic of the inversion (S1 Fig). This assay does not require cell viability. Strain SAM698 contains the inversion {P(on)} in all cells and was used as a standard for qPCR.

### Mating Assays

Cells were grown in 25 ml flasks in LB medium in a water bath at 37°C and 250 RPM shaking until they reached an OD_600_ of ~0.12. Arabinose and xylose (1% each) were added to donor strains to induce expression of P*xyl-rapI*, thereby causing cleavage of the ICE*Bs1* repressor ImmR [18,22] and de-repression of ICE*Bs1* gene expression. Where appropriate IPTG (2 μM) was added to induce expression of genes needed for complementation. Cultures were grown for 40 minutes after de-repression of P*xyl-rapI* to an OD600 of ~0.3-0.45.

Donors and recipients (in mid-exponential growth) were mixed at ratio of 2:1 (resulting in the equivalent of 3 OD600 units), vacuum-filtered, and filters were transferred to Spizizen minimal salts plates as described previously [17]. We found that these conditions resulted in high conjugation efficiencies that enabled detection of ICE*Bs1* integration by qPCR. Mating reactions were allowed to go for various amounts of time, as indicated, and were halted by placing filters in 3 ml of Spizizen minimal salts and vigorously vortexing. An aliquot of this mating mix was then used for serial dilutions and plating for colony forming units with appropriate antibiotics for selection of donors, recipients, and trasnsconjugants. The bulk of the mating mix was harvested for extraction of genomic DNA.

### Quantitative PCR assays to measure integration of ICE*Bs1* and Cre-mediated recombination

Quantitative PCR was used to measure integration of ICE*Bs1* in transconjugants. Cells were lysed with 40 mg/ml of lysozyme and genomic DNA was isolated using Qiagen DNeasy kit. qPCR was done using SSoAdvanced SYBR master mix and CFX96 Touch Real-Time PCR detection system (Bio-Rad).

The relative number of recipients (including transconjugants) in the mating mix was determined using primers to *mls* (*ermB*) that is only present in the recipient population (Δ*yddS*::*mls*). Primers to *mls* were oSAM403 (5’-GAAGGATTCT ACAAGCGTAC C-3’) and oSAM404 (5’-CTGGAACATC TGTGGTATGG -3’).

Integration in transconjugants was detected by the presence of the right junction (*attR*) at the endogenous location of ICE*Bs1* in *tnrS-leu2* (Fig 1B) using primers oSAM390 (5’- GCAAGTCTTCTCCCATAGC-3’) and oSAM399 (5’- GGCTTTTGTA AATAAAGATA TGATTTTACT AGGTTG -3’). Cre-mediated recombination (S1 Fig) was quantified using primers oSAM776 (5’- CCAGTCACGT TACGTTATTA GTTATAG -3’) and oSAM777 (5’- TACCGCACAG ATGCGTAAG -3’). Standard curves for *attR* and *mls* were generated using uninduced genomic DNA from SAM137 (mating experiments) or SAM698 (Cre experiments), ICE*Bs1*+ strains with copies of the relevant markers.

Serial dilutions were prepared from genomic DNA for each sample. The averages of at least three dilutions within the linear range were used to determine relative amount of ICE*Bs1* integration or Cre-mediated recombination. These were normalized to CFUs, as indicated, to determine the fraction of the recipient or transconjugant population with each marker.

### Microscopy and image acquisition

We used an inverted microscope (Nikon, Ti-E) with a motorized stage placed in a temperature-controlled box (In Vivo Scientific) at 37°C. Images were acquired with a CoolSnap HQ2 camera (Photometrics). Fluorescence was generated using a Nikon Intensilight mercury illuminator through appropriate sets of excitation and emission filters (filter set 49000 for DAPI, 49003 for mNeongreen, and 49008 for mApple; Chroma). Samples were spotted onto agarose pads that were set in a homemade incubation chamber made by stacking two sealable Frame-Seal Incubation Chambers (BIO-RAD). Agarose pads contained 1.5% Ultrapure agarose (Thermo Fisher) dissolved in either minimal medium with Spizizen salts or CH medium [54]. Minimal medium was used when assessing fluorescence intensity and CH medium was used for tracking cell viability. For time-lapse experiments, agarose pads were supplemented with the appropriate carbon source at 1% L-arabinose, 1% glucose, and 1% D-xylose w/v. Where applicable, agarose pads were supplemented with 25 ng/ml DAPI, 170 ng/ml FM4-64, and 0.1M propidium iodide. For visualization of cell membranes, cultures were incubated with FM4-64 (170 ng/ml) for 1 hour before imaging.

### Microscopy image analysis

Images and videos were processed with Fiji (ImageJ) [67]. All images for fluorescent channels were subjected to background subtraction using a rolling ball radius of 10 or 20 pixels (10 pixels was used for images with foci). Cell meshes were acquired using Oufti [68]. Meshes were analyzed in MATLAB using custom scripts.

## Acknowledgments

We thank Mary Anderson and Alam Garcia Heredia for comments on the manuscript and Emily L. Bean for the reporter system for detecting Cre-mediated recombination.

## Supporting information

**S1 Fig.**
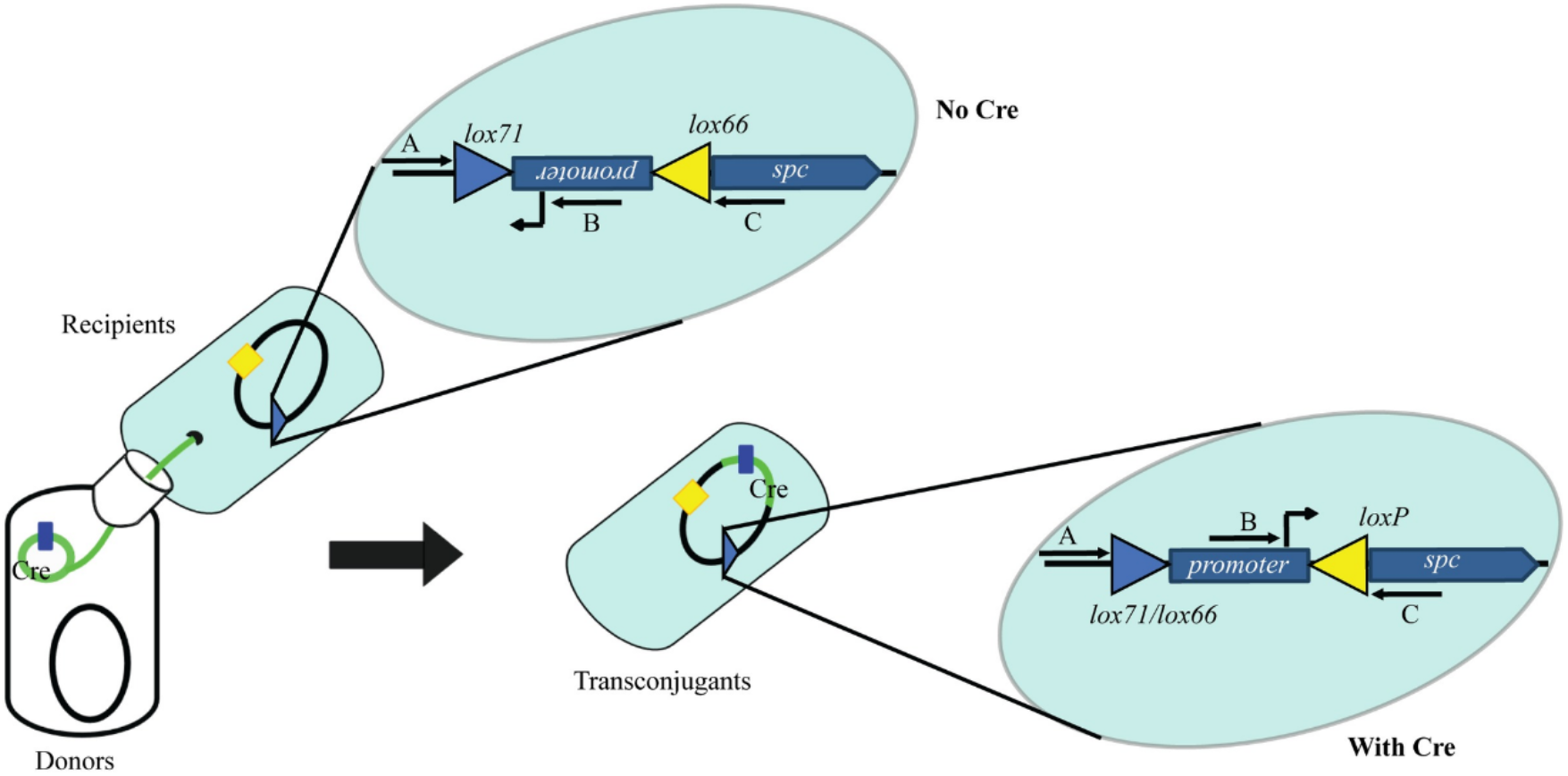
Conceptual schematic for mobilizable Cre recombinase assay to measure transfer of ICE*Bs1*. Donors containing ICE*Bs1* with P*xis-cre* were mated with recipients harboring a reporter to indicate Cre-mediated recombination (inversion). Transfer of ICE*Bs1* with P*xis-cre* and production of Cre results in inversion of the DNA fragment that can be detected by qPCR. The green circle in the cartoon at the bottom left represents ICE*Bs1* that expresses *cre*. The blue triangle in recipients and transconjugants indicates the reporter for and result of Cre-mediated inversion. Blue and yellow triangles indicate *lox* sites as labeled. The region between the blue and yellow triangles contains a promoter, either driving transcription of *spc* (With Cre; bottom right) or in the opposite orientation (No Cre; top). Blue block arrow represents *spc* open reading frame. Inversion in the presence of Cre produces *lox71/lox66* and *loxP* sites which are incapable of subsequent recombination [61].The horizontal black arrows with letters A, B, and C represent primers for PCR to detect the reporter and Cre-mediated inversion. PCR with primers B and C detect the product of Cre-mediated recombination. PCR with primers B and A detect the starting reporter with the promoter in the ‘off’ orientation.

**S2 Fig.**
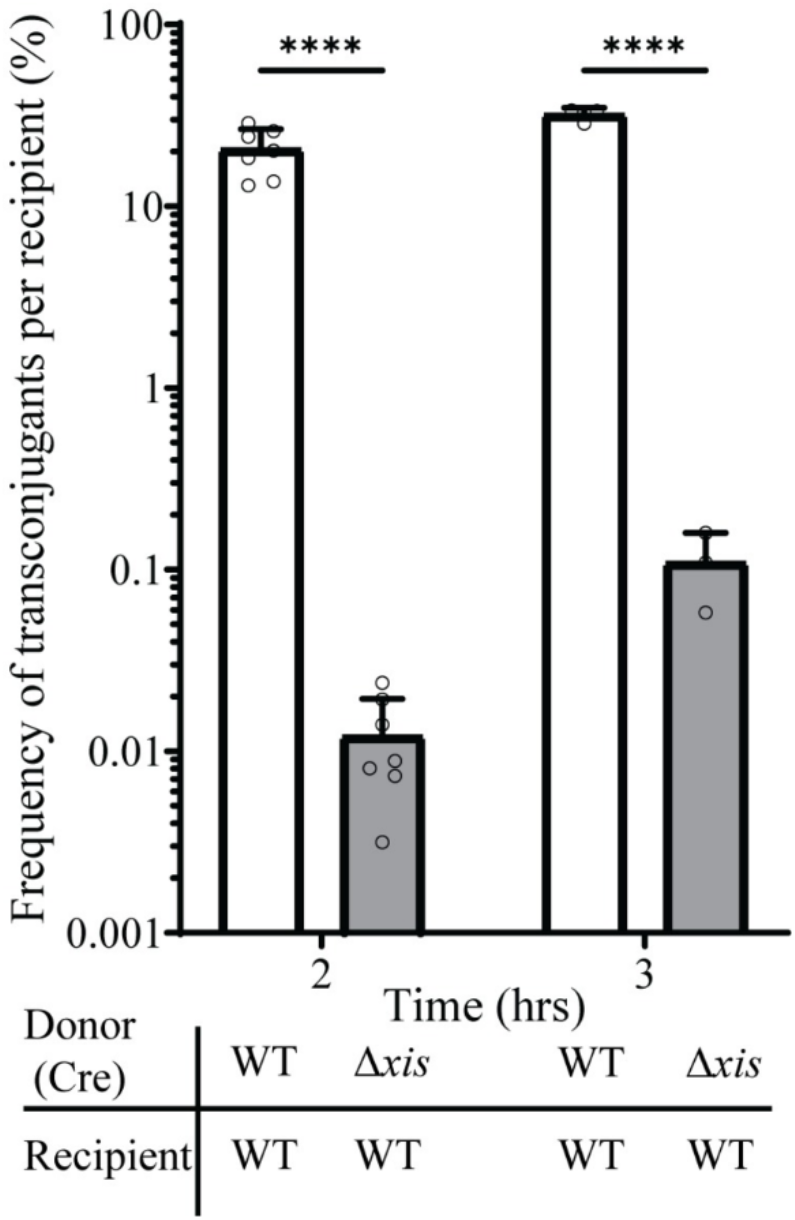
Acquisition efficiency per recipient as measured by CFUs. Acquisition of ICE*Bs1* by recipients was measured by selective plating and normalized to the total number of recipients. The identity of donor and recipient strain is indicated below the x-axis. Donors: WT (SAM830); Δ*xis* donor (SAM892). Recipients: WT (SAM599). Results are from at least 3 biological replicates and error bars indicate standard deviation. The differences between the wild type and Δ*xis* are statistically significant at both two and three hours. Statistical comparison: t-test, **** P value < 0.00005.

**S3 Fig.**
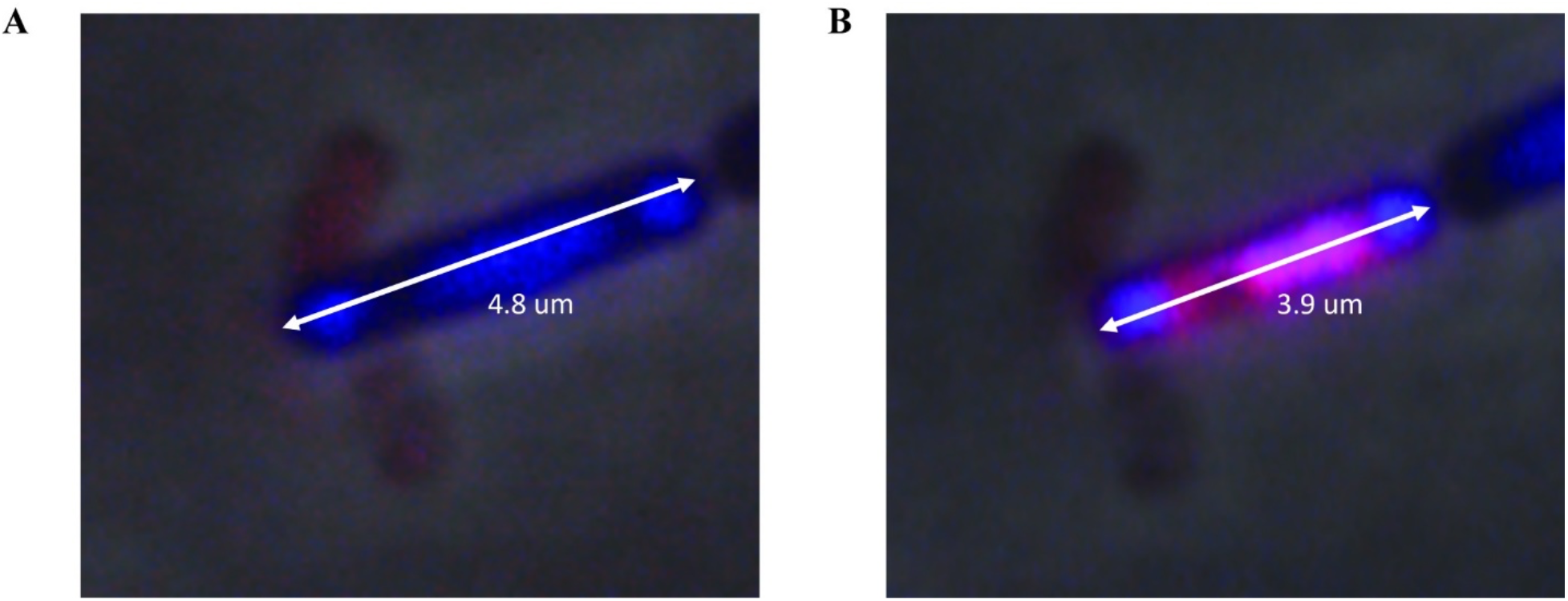
Cell death precedes cell lysis. Cells were (ICE*Bs1* Δ*xis*; SAM427) grown at 37°C in defined minimal medium with L-arabinose. Expression of *rapI* (P*xyl-rapI*) was induced by the addition of D-xylose. After 60 mins, cells were washed and spotted onto agarose pads containing CH medium, DAPI, and propidium iodide. **A).** Cell 3 mins prior to PI uptake. The white arrows indicate the axis across which cell length was measured and the cell length is indicated under the arrow. DAPI is pseudo-colored blue. Propidium iodide is pseudo-colored red. **B)** Cell from **A**, at the time of uptake of propidium iodide and 3 mins prior to lysis.

**S4 Fig.**
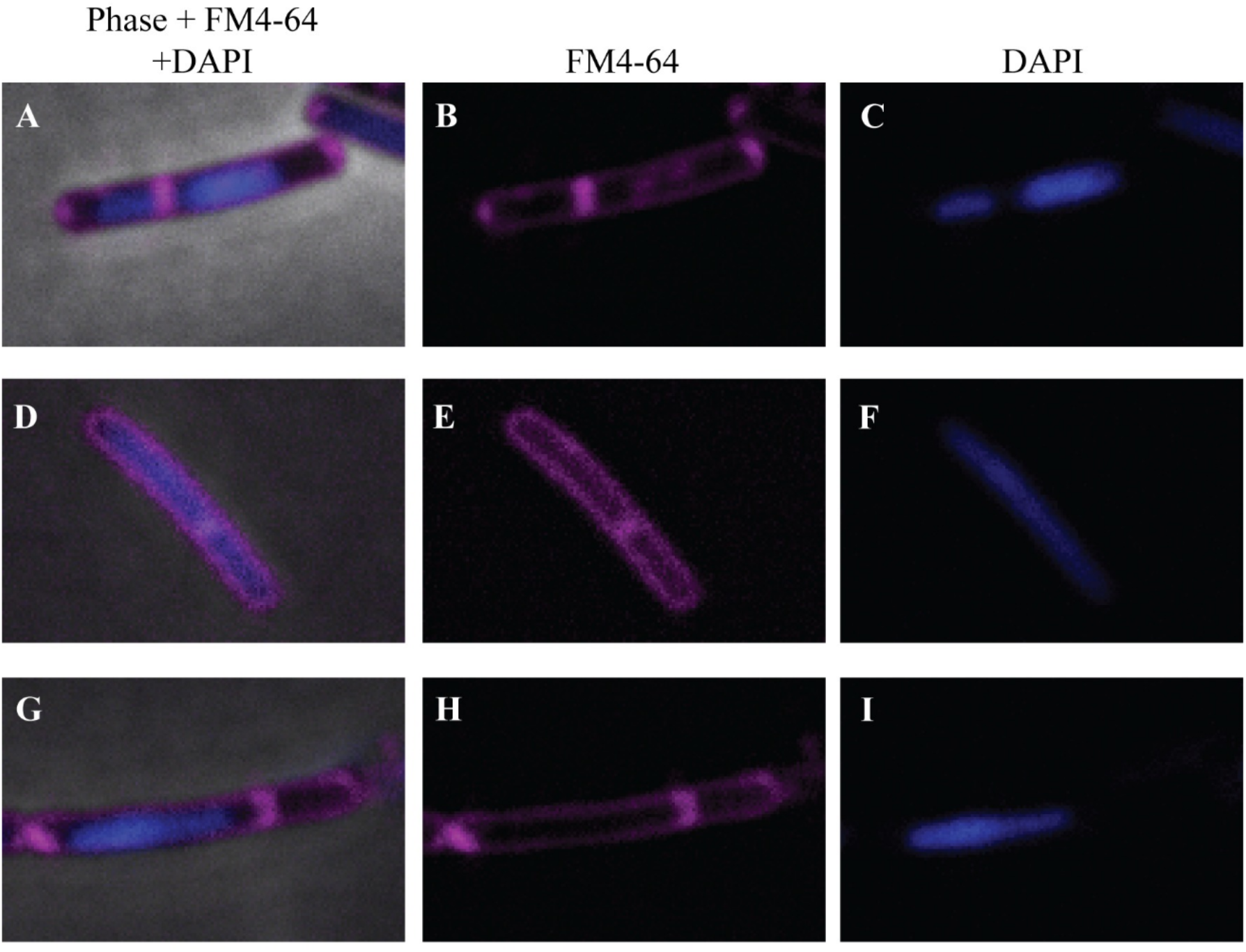
Activation of ICE*Bs1* Δ*xis* causes chromosome abnormalities and defects in cell division. Cells (ICE*Bs1 Δxis;* SAM427) were grown at 37°C in defined minimal medium with L-arabinose. Expression of *rapI* (P*xyl-rapI*) was induced by the addition of D-xylose. After 60 mins, cells were washed and spotted onto agarose pads containing CH medium, DAPI, and FM4-64. **A-C)** Cell with asymmetric chromosome segregation and mislocalized septum formation. **D-F)** Cell with an elongated nucleoid and a guillotined chromosome. **G-I)** Cell with asymmetric chromosome segregation and anucleate division. **A,D,** and **G**) Composite image of phase, FM4-64 and DAPI channels. **B,E,** and **H)** FM4-64 channel. **C,F,** and **I)** DAPI channel.

**S5 Fig.**
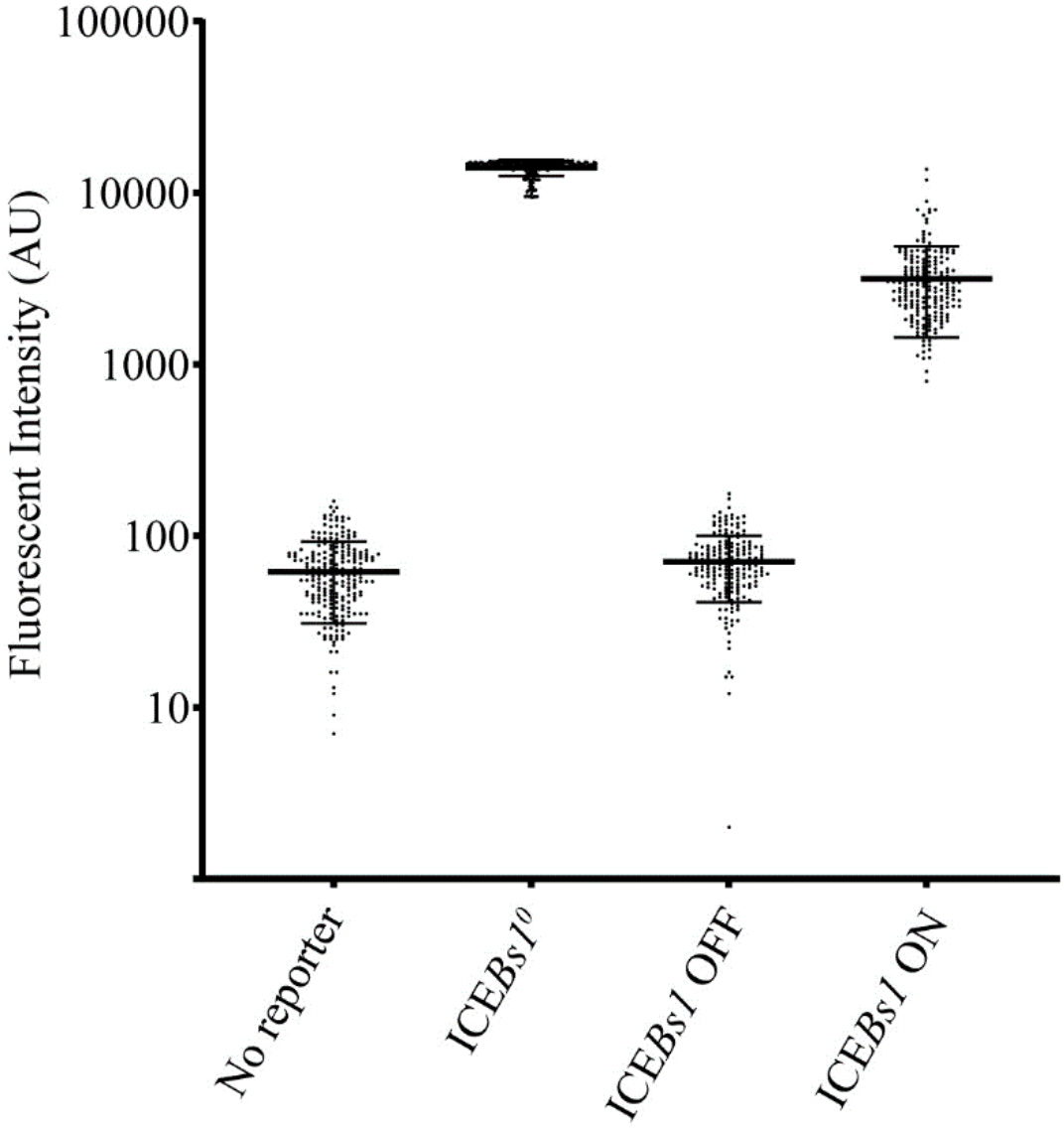
Expression from P*xis* in cells with and without ICE*Bs1*. Cells with the reporter P*xis-gfp* were grown at 37°C in a defined minimal medium with L-arabinose. Expression of *rapI* (P*<u>xyl</u>-rapI*) was induced by the addition of D-xylose. After 2 hrs cells were spotted onto agarose pads and imaged for fluorescence. Each data point represents the fluorescent intensity from a single cell. the longest horizontal line represents the mean and bars above and below represent the standard deviation. No reporter is the parental strain with no genetic markers (AG174). ICE*Bs*1^0^ is a strain cured of ICE*Bs1* (SAM011) and expression from P*xis* is in the absence of the repressor ImmR. ICE*Bs1* OFF is a strain containing ICE*Bs1* Δ(*rapI-phrI*)::*kan* (SAM032) without exogenous activation of the element. These cells produce active ImmR and transcription from P*xis* is repressed and there is only basal expression of ICE*Bs1*. ICE*Bs1* ON is the same strain (SAM032) except that ICE has been activated by virtue of expression of *rapI* and subsequent inactivation of the repressor ImmR. P*xis* is expressed under these conditions.

**S6 Fig.**
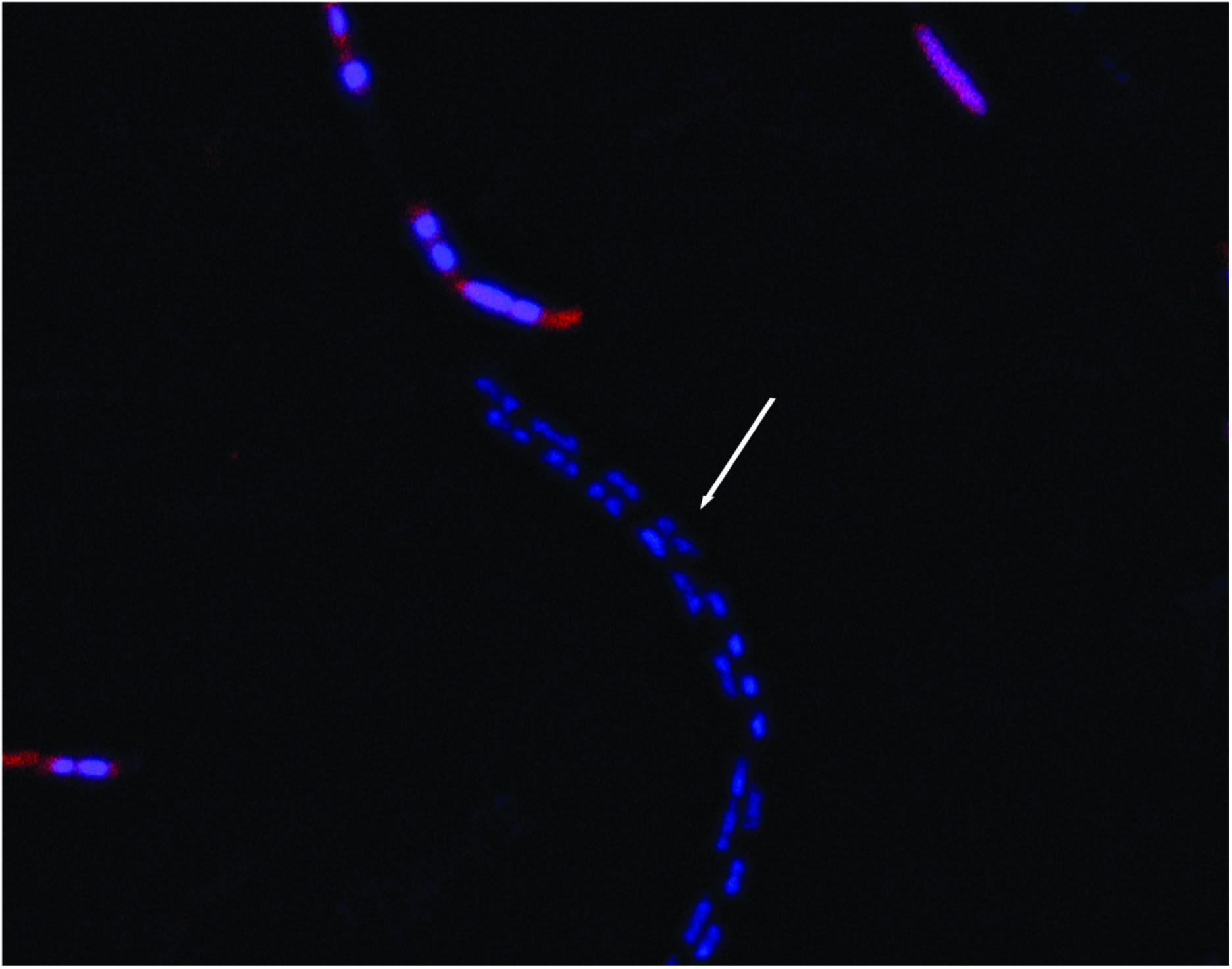
Cells that continue to grow normally following activation of ICE*Bs1* Δ*xis* in the bulk culture still have the element. Cells containing ICE*Bs1* Δ*xis* and the reporter P*xis-mApple*(SAM456) were grown at 37°C in a defined minimal medium with L-arabinose. Expression of *rapI* (P*xyl-rapI*) was induced by the addition of D-xylose. After 60 mins cells were washed and spotted onto agarose pads with CH medium and DAPI. The image shown is after 3 hrs on the agarose pad and is a composite of the red fluorescence (P*xis-mApple*) and DAPI channels. Red fluorescence is pseudo-colored red and DAPI is pseudo-colored blue. The white arrow points to chain of normally growing cells that have ICE*Bs1* but the element has not been activated. Had they lost ICE*Bs1*, or had ICE*Bs1* been activated, then P*xis-mApple* would be expressed.

**S1 Movie. The fate of ICE*Bs1* Δ*xis* transconjugants**. Donors (ICE*Bs1* Δ*xis*, SAM472) harboring ICE*Bs1* with constitutively expressed mNeongreen were mated with recipients (SAM271) constitutively expressing mApple for 4 h. Following mating, samples were spotted onto nutrient-rich agarose pads containing erythromycin and lincomycin (recipients and transconjugants contain *mls* and are resistant to these two antibiotics). Donors appear pale yellow. Recipients appear red. Transconjugants are both red and yellow and appear orange or bright yellow.

**S2 Movie. Activation of ICE*Bs1* Δ*xis* (unable to excise) causes cell lysis.** Cells (SAM427) were grown at 37°C in a defined minimal medium with L-arabinose. Expression of *rapI* (P*xyl-rapI*) was induced by the addition of D-xylose. After 60 mins cells were washed and spotted onto to nutrient-rich agarose pads and imaged every 3 mins for 3 hrs.

**S3 Movie. Activation of WT ICE*Bs1* does not cause cell lysis.** Cells (SAM426) were grown at 37°C in a defined minimal medium with L-arabinose. Expression of *rapI* (P*xyl-rapI*)was induced by the addition of D-xylose. After 60 mins cells were washed and spotted onto nutrient-rich agarose pads and imaged every 3 mins for 3 hrs.

**S4 Movie. Activation of ICE*Bs1* unable to excise causes chromosome abnormalities.** Cells (SAM427) were grown at 37°C in a defined minimal medium with L-arabinose. Expression of *rapI* (P*xyl-rapI*) was induced by the addition of D-xylose. After 60 mins, cells were washed and seeded to nutrient-rich agarose pads containing DAPI and imaged every 15 mins for 45 mins and then every 3 mins for 2.25 hrs.

**S5 Movie. Chromosome abnormalities caused by activation of ICE*Bs1* that is unable to excise are not due to SPβ, PBSX, or *skin*.** Cells (SAM529) were grown at 37°C in a defined minimal medium with L-arabinose. Expression of *rapI* (P*xyl-rapI*) was induced by the addition of D-xylose. After 60 mins cells were washed and spotted onto nutrient-rich agarose pads containing DAPI and imaged every 15 mins for 45 mins and then every 3 mins for 2.25 hrs.

**S6 Movie. Cells containing ICE*Bs1* Δ*xis* that survive induction are those that have not activated the element.** Cells (SAM456) were grown at 37°C in a defined minimal medium with L-arabinose medium. Expression of *rapI* (P*xyl-rapI*) was induced by the addition of D-xylose. After 60 mins cells were washed and spotted onto nutrient-rich agarose pads containing DAPI and imaged every 15 mins for 45 mins and then every 3 mins for 2.25 hrs.

